# Pangenome analyses of tea plant reveal structural variations driven gene expression alterations and agronomic trait diversification

**DOI:** 10.1101/2025.03.14.643311

**Authors:** Lingling Tao, Junyan Zhu, Jianbing Hu, Jun Wu, ChunLin Chen, Youyong Li, Fangdong Li, Hongrong Chen, Songyan Huang, Qianqian Zhou, Yuanyan Zhao, Yanping Hu, Shengrui Liu, Kun Dong, Linbo Chen, Benying Liu, Xiaochun Wan, Enhua Xia, Yongfeng Zhou, Chaoling Wei

## Abstract

Tea plants, one of the world’s most economically significant beverage crops, exhibit extensive genetic diversity and a rich array of secondary metabolites. While structural variations (SVs) are key drivers of phenotypic innovation, their regulatory roles in transcriptional networks and agronomic trait diversification remain underexplored in this perennial crop. We constructed a pangenome from 21 representative tea accessions and their wild relatives. Comparative analysis revealed that wild relatives harbor a greater abundance of gibberellin-related genes, whereas cultivated accessions exhibit an expansion of defense-related genes. We identified 522,428 non-redundant SVs, primarily insertions/deletions (73.56%), which were enriched in promoter regions. These SVs influenced the expression of genes governing flavor compounds, including those involved in flavonoid, amino acid, and terpenoid biosynthesis. Population SV analysis of 275 tea accessions uncovered three haplotypes in the *ANS3* gene promoter region, with Hap1, containing a 192 bp insertion, predominantly found in wild relatives but largely lost in modern cultivars. This insertion enhanced *CtANS3* expression, increasing anthocyanin accumulation. Additionally, a 159 bp insertion in the *CtLRR1* promoter reduced resistance to *Colletotrichum gloeosporioides* in wild relatives. Our findings underscore SVs as pivotal regulators of genomic diversity, flavor differentiation, and adaptive evolution in tea plant domestication.

## Introduction

Tea is one of the world’s most valuable agricultural commodities, driving significant economic activity and playing a central role in international trade^1^. The two main cultivated tea varieties, *Camellia sinensis var. sinensis* (CSS) and *Camellia sinensis var. assamica* (CSA), are widely grown and contribute substantially to global tea production^2^. Additionally, *Camellia taliensis* (CT), a wild relative, is distributed primarily in southwestern China, and exhibits extensive genetic diversity^3,4^. Despite the need for continuous genetic improvement to meet evolving consumer preferences, the domestication of tea plants has reduced genetic diversity, creating bottlenecks that constrain breeding efforts^5^. Compared to cultivated accessions, wild relatives display higher genetic diversity^3^ and elevated phenolic acid content but lower flavan-3-ol levels ^6–8^. This narrowing of genetic variation necessitates a deeper exploration of wild tea germplasm to enhance breeding programs.

Structural variations (SVs), including insertions, deletions, inversions, and translocations, play critical roles in plant evolution, domestication, and breeding, influencing key traits such as flowering time^9,10^, fruit flavor^11^, stress resistance^12^, and environmental adaptability^13^. Recent studies have highlighted the impact of SVs on agronomic traits across various crops. For example, Wang et al. discovered that a 209 bp non-coding RNA insertion within the intron of *MMK2*, a mitogen-activated protein kinase homolog, regulates fruit coloration in apple via SV-mediated expression changes^14^. Similarly, in soybean, a 10 kb presence-absence variation was identified as a key determinant of seed luster variation ^15^. These findings underscore the significance of SVs in crop improvement and adaptation.

However, traditional reference-genome-based analyses struggle to fully capture SV diversity, leading to reference bias. While whole-genome resequencing effectively identifies SNPs and small indels, detecting SVs using short-read sequencing remains technically challenging^16^. The advent of high-throughput long-read sequencing has significantly improved SV detection across crops^17^. For instance, *de novo* assembly of 12 maize inbred genomes revealed extensive SVs that contributed to heterosis^18^, while comparative genomic analysis in tomato identified beneficial SV-associated genes from wild relatives, enabling crop improvement^19^.

Despite recent advancements in tea plant pangenome research, existing studies remain limited. Prior efforts constructed pangenomes using Continuous Long Reads (CLR) from 22 tea accessions^20^, but these focused primarily on cultivated varieties and lacked systematic SV analyses of wild relatives. Additionally, most genome assemblies relied on Continuous Long Read (CLR) technology, limiting resolution for detecting complex SVs such as large insertions, deletions, and inversions. Current pangenomes also lack saturation, with inadequate representation of wild relatives from southwestern China and CSA accessions. Future research should integrate high-quality genomes with multi-omics approaches to comprehensively characterize SVs and their functional impact, providing valuable genetic resources for tea breeding.

To address these gaps, we assembled five newly cultivated tea genomes and one wild relative genome using long-read sequencing, integrating them with 15 published genomes^3,5,8,20,22,23^. This enabled the construction of a comprehensive pangenome variation map. By incorporating transcriptomic data, we identified numerous non-redundant SVs and demonstrated how large insertions and deletions in promoter regions regulate gene expression and phenotypic differentiation. Population-level SV analysis of 275 accessions revealed a 192 bp insertion in the *CtANS3* promoter that enhanced anthocyanin accumulation in wild relatives. Additionally, a 159 bp insertion in the *CtLRR1* promoter affected disease resistance. These findings highlight SVs as key regulators of gene expression and suggest that their enrichment in promoter regions drives phenotypic diversity during domestication. Our study provides a theoretical foundation for future genetic improvement of tea plants.

## Results

### High-quality genome assembly of representative tea plants

The genome assembly of 21 tea accessions, including two wild relatives (*Camellia taliensis*, CT) and 19 cultivated accessions (seven *Camellia sinensis var. assamica*, CSA; one *Camellia sinensis var. pubilimba*, CSP; and eleven *Camellia sinensis var. sinensis*, CSS), was analyzed to capture the genetic diversity of tea plants (Table 1). High-quality chromosome-scale reference genomes were generated for five cultivated accessions (three CSA: MHDY, QS3, YK37; and two CSS: HJY, ZC102) using PacBio CCS technology, with sequencing depths of 24.19–60.92× (Table 1; Supplementary Table 1). Additionally, a novel genome for the wild relative DLC was assembled, forming the foundation for comparative genomic analyses. These five HiFi-based assemblies averaged 3.18 Gb in size, with contig N50 values ranging from 131.14 to 142.81 Mb. An average of 96.46% of the assembled contigs were successfully anchored to 15 pseudochromosomes (Table 1). Assembly completeness was validated using Benchmarking Universal Single-Copy Orthologs (BUSCO), yielding scores of 98.82– 99.32%. Furthermore, the long terminal repeat (LTR) assembly index (LAI) ranged from 12.59 to 14.74, meeting high-quality reference genome standards. To construct haplotype-resolved assemblies, Hifiasm was used to integrate HiFi reads with Hi-C data, generating both haploid1 and haploid2 assemblies. The final 10 chromosome-scale haplotype-resolved sequences were aligned to the reference genome using RagTag, achieving an average contig N50 of 209 Mb, with a mean GC content of 39.1% and a BUSCO completeness score of 98.5% (Supplementary Table 3).

**Table 1.**
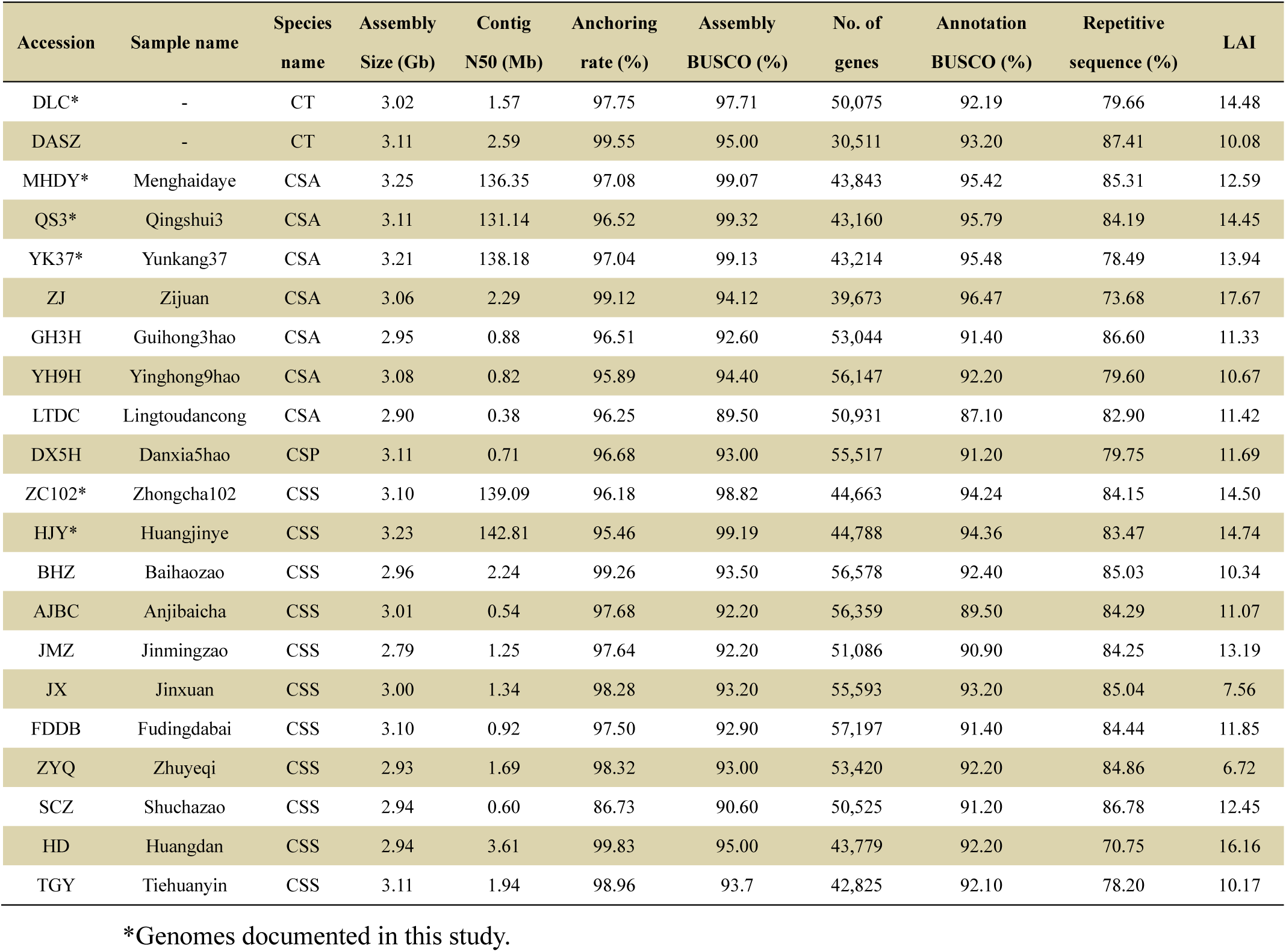
Summary of the Assembly and Annotation of 21 Tea Plant Genomes.

Gene annotation combined homology-based predictions with transcriptome data (Methods), identifying 43,160–44,788 protein-coding genes per genome, with an average BUSCO completeness score of 95.06%. Phylogenetic analysis clustered cultivated and wild tea plants into two major clades, with CSA and CSS further diverging into distinct subclades, while CSP grouped closely with CSS. This phylogenetic structure aligns with previous reports on tea population genetics^23^. Divergence time estimates indicated that wild and cultivated tea plants split approximately 5.84 million years ago (MYA) (Fig. 1a).

**Fig. 1.**
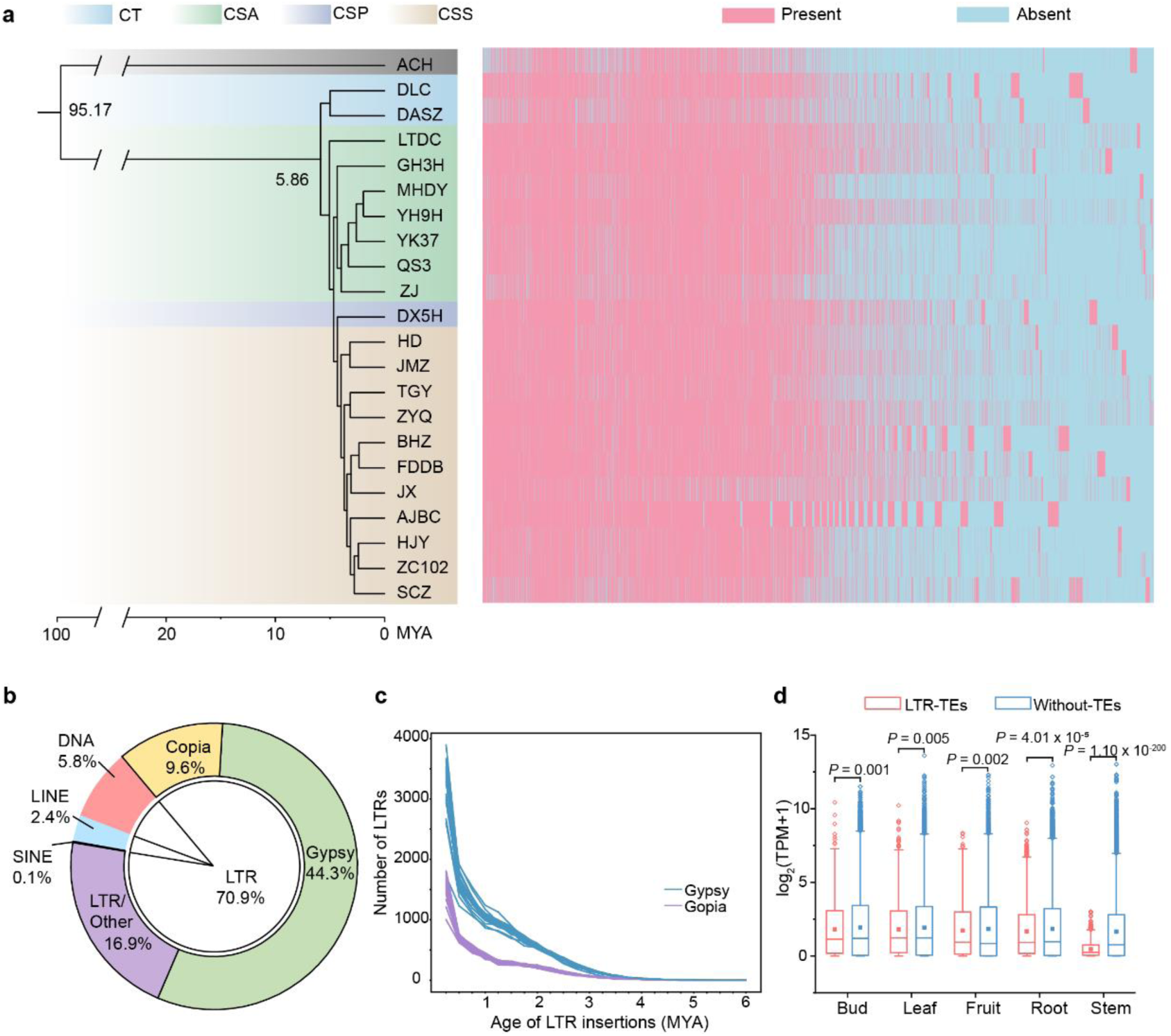
| Phylogenetic analysis and transposable element dynamics in tea plant genomes. **a**, Phylogenetic analysis of 21 tea plant genomes using kiwifruit as an outgroup, with presence/absence variations in the pan-gene family. The genomes cluster into four clades: wild relative CT and cultivated groups (CSA, CSS, CSP). **b,** LTR content proportions in the genomes assembled in this study, with Gypsy elements (44.3%) predominating over Copia (9.6%). **c,** Insertion time distribution of full-length Copia and Gypsy LTRs across the 21 genomes. Each line represents an individual genome from panel (**a**), with Copia elements in blue and Gypsy in purple. **d,** Expression levels of genes with LTR insertions (n = 2,621) versus genes without TE insertions (n = 36,654) across various tissues of *C. sinensis* cv. Zhongcha102. Expression levels were measured in buds, leaves, fruits, roots, and stems. Box plots indicate median expression values, and statistical significance was assessed using a two-tailed Student’s *t*-test. TPM: transcripts per million mapped reads.

Repetitive sequences accounted for 70.75–87.45% of the assembled genomes, with LTR retrotransposons (LTR-RTs) constituting the dominant fraction. Across the newly assembled genomes, LTR-RTs made up an average of 70.9% of the genomic content (Fig. 1b). Analysis using LTR_retriever identified 17,388–25,566 full-length LTR-RTs (fl-LTRs) per genome (Supplementary Table 5). A comparative analysis of LTR insertion times revealed that Gypsy and Copia elements underwent continuous expansion approximately 4 MYA, followed by a pronounced burst around 0.25 MYA. Notably, Gypsy elements exhibited significantly higher insertion activity than Copia elements, suggesting more extensive expansion of the Gypsy family (Fig. 1c). LTR-RTs predominantly accumulated in intergenic regions (Supplementary Fig. 1), with an average of 2,148 genes per genome affected by LTR insertions (Supplementary Table 5). Transcriptomic analysis across various tissues of ZC102 revealed that genes lacking LTR insertions exhibited significantly higher expression levels than those with LTR insertions (Fig. 1d).

### Gene gain and loss during tea plants domestication

We constructed a gene-based pangenome across 21 genomes using orthofinder clustering on the annotated gene models. The number of pan gene families rapidly to a platform with the increasing genome numbers, while the number of core gene families showed a similar decreasing trend, indicating the representativeness of our pangenome (Fig. 2a). A total of 8,317 core gene families were shared across all 21 genomes, constituting 18.7% of the total clusters (Supplementary Fig. 2a), while the genes within these core families represented 46.1% of all genes (Supplementary Fig. 2b). Softcore gene families, defined as those present in 19 or 20 genomes, comprised 20.5% of the total clusters. Dispensable gene families, found in 2 to 18 genomes, and private gene families, present in only one genome. The coding sequence (CDS) length and expression levels of the core genes were higher compared to the softcore and dispensable genes, but lower Ka/Ks ratios may suggest the functional conservation of the former (Supplementary Fig. 2c-e).

**Fig. 2.**
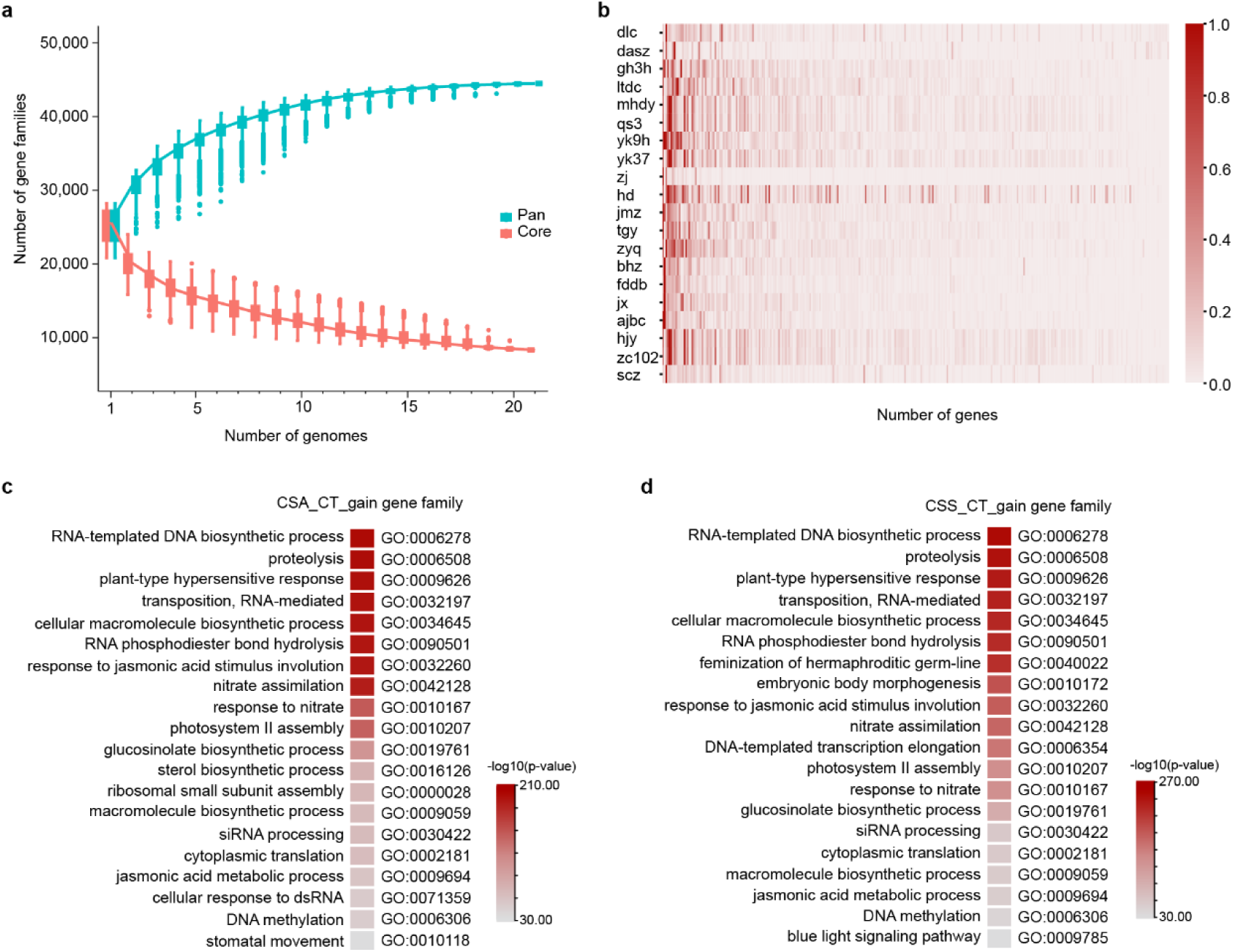
| Pangenome analysis and gene gain/loss patterns. **a**, Variation of gene families in pangenome and core genome. **b**, Gene information of gene families across 21 accessions is highly abundant. The color indicates the ratio of the gene content in one accession to the maximum gene content within that gene family. **c**, GO enrichment analysis of genes gained in the CSA population, compared with CT. **d**, GO enrichment analysis of genes gained in the CSS population, compared with CT.

To investigate gene content variation among tea accessions, we analyzed the pangenome across CT, CSA, and CSS populations. GO enrichment analysis revealed significant overrepresentation of 45 and 24 gene families in the CT population compared to CSA and CSS, respectively (Mann–Whitney–Wilcoxon test, *P* < 0.05; Supplementary Table 6). These genes were predominantly associated with gibberellin and auxin signaling pathways, as well as cell division processes, which likely reflects the tall, tree-like growth habit of wild relatives (Supplementary Fig. 3a-b). In contrast, CSA and CSS exhibited enrichment in 153 and 223 gene families, respectively, with a notable concentration in defense-related pathways, including those involved in disease and insect resistance (*e.g*., jasmonic acid metabolism) (Fig. 2c-d). The CSS population also displayed a significant expansion in 70 gene families relative to CSA, with enrichment in pathways related to temperature adaptation, pollen tube development, protein trafficking, and gravitropic responses (Supplementary Fig. 3c-d), highlighting distinct selection pressures during domestication.

### Extensive SVs within 21 tea plant genomes

To identify large-scale SVs in the tea plant pangenome, we aligned 20 genome assemblies to the reference genome (ZC102) and used SyRI software to classify SVs into four categories: inversions, translocations, insertions, and deletions (>50 bp). A total of 522,428 non-redundant SVs were identified (Supplementary Table 7). SV numbers were significantly lower in the DLC and DASZ genomes compared to cultivated tea plants, indicating distinct genomic characteristics between wild and domesticated populations (Fig. 3a). Insertions and deletions were the most prevalent SV class, comprising 384,312 events (73.56% of total SVs), followed by translocations (25.07%), while inversions were the least frequent (1.37%) (Supplementary Table 7). The predominance of large-scale insertions and deletions suggests their central role in shaping genomic variation in tea plants (Fig. 3a).

**Fig. 3.**
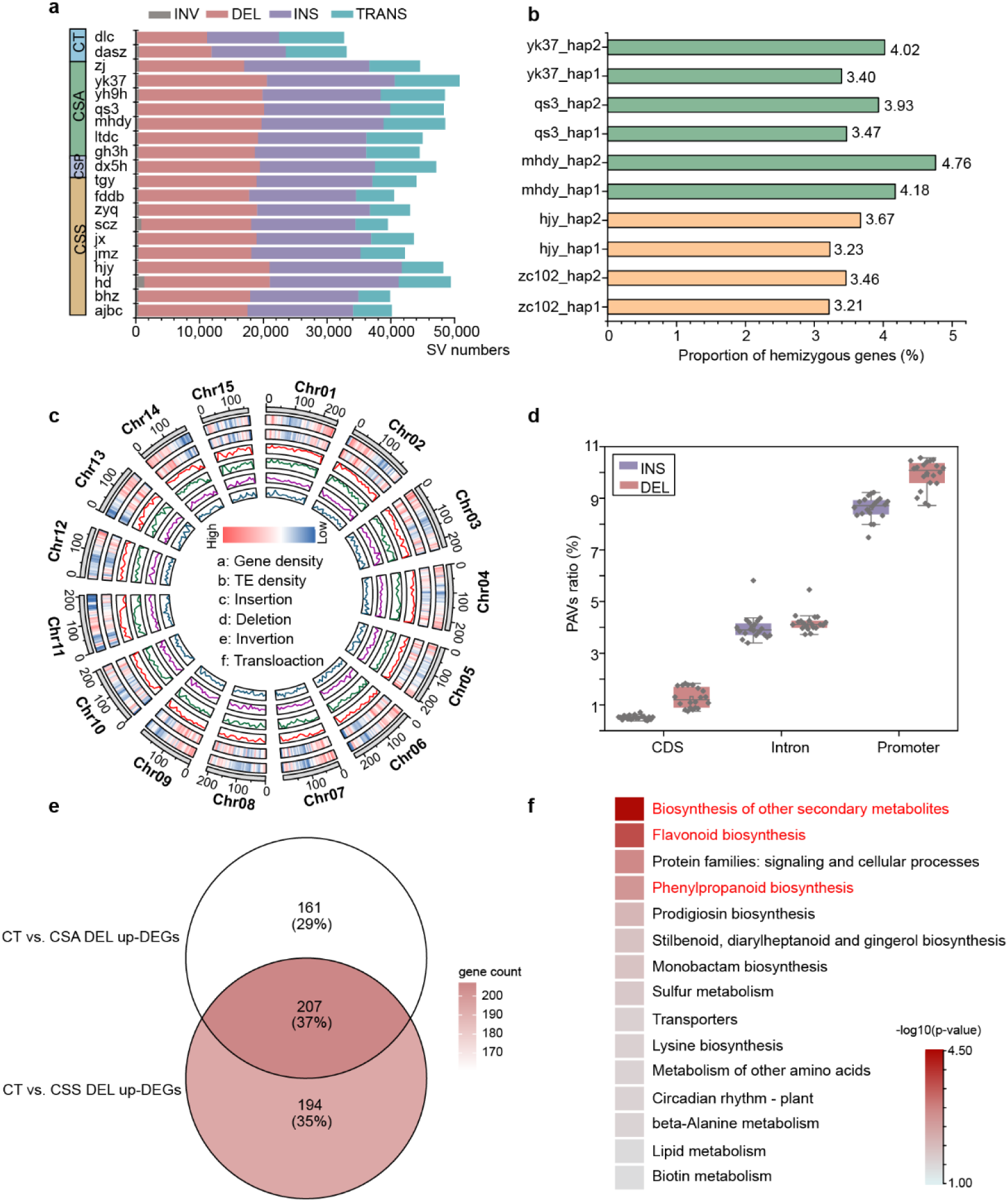
| Structural variations in the tea plant genomes. **a**, Distribution of four SV types across tea plant accessions, with colors indicating population membership (CT, CSA, CSS, CSP). **b**, Proportion of hemizygous genes in each haplotype genome, with green representing CSA accessions and yellow representing CSS accessions. **c**, SV distribution across 15 chromosomes in 20 genomes. **d**, Percentage overlap between SVs and different genomic regions across 20 genomes, with red dots representing deletions and purple dots representing insertions. **e**, Intersection of upregulated differentially expressed genes between CT *vs*. CSA and CT *vs*. CSS, highlighting genes with promoter-region deletions based on transcriptome analysis. **f**, KEGG enrichment of genes identified in panel **e**.

The application of long-read sequencing has greatly improved SV detection, enabling a more comprehensive characterization of hemizygous genes resulting from structural rearrangements in heterozygous diploid genomes. We analyzed ten haplotype genomes, defining hemizygous genes as those flanked by heterozygous insertion/deletion variants. The proportion of hemizygous genes ranged from 3.21% to 4.76%, with CSA accessions exhibiting a mean frequency of 3.96% (n = 4), slightly higher than the 3.39% observed in CSS accessions (n = 6) (Fig. 3b). GO enrichment analysis showed that hemizygous genes were primarily associated with plant-type primary cell wall biogenesis, cellulose biosynthesis, and ovule development, suggesting a role in male gametophyte development (Supplementary Fig. 4).

Consistent with previous findings, our analysis confirmed the substantial transposable element (TE) content in the tea plant genome. Insertions and deletions were enriched in gene-dense regions, displaying similar spatial distribution patterns. In contrast, inversions and translocations were significantly less frequent and exhibited a random chromosomal distribution (Fig. 3c).

### SVs influence the differential expression of flavor-related genes among tea accessions

SVs can modulate gene expression by altering gene sequences or modifying cis-regulatory elements ^24^. Since insertions and deletions were the predominant SV types identified in our analysis, we examined their genomic distribution to assess their functional impact. By integrating variant location data with genomic features (CDS, introns, and promoter regions), we found a notable enrichment of insertions and deletions in promoter regions relative to coding and intronic regions. Specifically, insertion and deletion frequencies in promoter regions reached 8.62% and 9.91%, respectively (Fig. 3d). Given the essential role of promoters in regulating gene expression, we performed transcriptome analyses on an expanded dataset of 21 tea accessions, evenly divided into three groups (CT, CSA, CSS; n = 7 per group). Differential expression analysis of genes harboring promoter deletions revealed 207 commonly upregulated genes significantly enriched in the flavonoid biosynthesis pathway and 397 commonly downregulated genes enriched in amino acid metabolism (Fig. 3e-f; Supplementary Fig. 5a). Similarly, for genes with promoter insertions, we identified 216 commonly upregulated genes between wild and cultivated tea plants, predominantly involved in terpenoid biosynthesis, suggesting a potential role in modulating tea aroma via volatile compound synthesis (Supplementary Fig. 5b-c). These findings highlight that promoter insertions and deletions play a crucial role in regulating genes associated with flavonoid, amino acid, and terpenoid metabolism in tea plants.

### A 192 bp insertion in the *CtANS3* promoter enhances anthocyanin accumulation

Despite increasing interest in tea genomics, large-scale SV analyses across diverse tea populations remain limited. To address this gap, we constructed a pangenome variation map by integrating variant data from multiple genomes, anchoring it to the ZC102 reference genome with alternative allelic sequences. Using vg call, we genotyped SVs across 275 tea accessions (Supplementary Table 8). To identify domestication-related selection signatures, we analyzed genomic regions under selective pressure during the transition from wild to cultivated tea. Among the top 5% *F*_st_ selection sweep regions differentiating CT and CSS populations, we identified key genes related to cold response and disease resistance, including *COR* (cold-regulated protein), two tandemly duplicated *RPV* (disease resistance) genes, and *OFP* (ovate family proteins). In the CSA-to-CSS domestication transition, we observed selection on genes influencing tea quality, such as *HCT* (shikimate O-hydroxycinnamoyltransferase), *CHI* (chalcone isomerase), and *LAR* (leucoanthocyanidin reductase), all involved in flavonoid biosynthesis, as well as *TPS* (terpene synthase), which regulates aroma compound synthesis (Fig. 4a). Notably, an LAR-associated 184 bp SV defined two haplotypes (Hap1/Hap2) with strong population-specific differentiation (Supplementary Fig. 6a). While 82% of CT accessions carried Hap2 (compared to only 3% Hap1), domesticated populations exhibited a marked shift toward Hap1 (CSA: 51%, CSS: 37%) (Supplementary Fig. 6b). Transcriptome analysis confirmed significantly lower *LAR* expression in CT relative to domesticated accessions (*P* < 0.01) (Supplementary Fig. 6c), suggesting positive selection of Hap1 for enhanced expression during domestication.

**Fig. 4.**
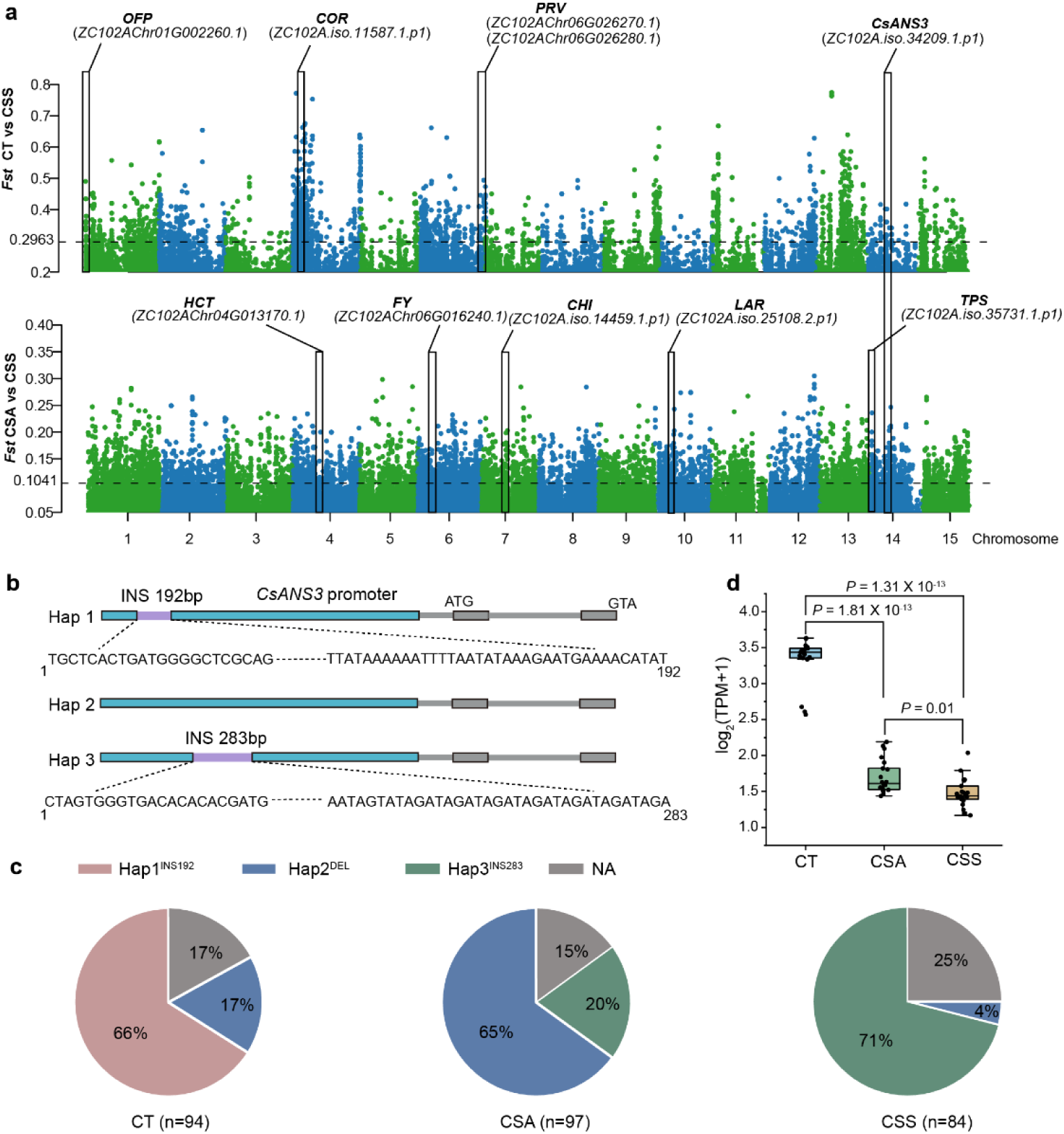
| Selection signatures in tea populations based on SVs. **a**, *F*_ST_ distribution for SVs between wild relatives (CT) and cultivated tea populations (CSA, CSS), with the black dashed line indicating the top 5% threshold. **b**, Schematic representation of three haplotypes in the *ANS3* gene structure. **c**, Frequency distribution of three *ANS3* haplotypes (Hap1^INS192^, Hap2^DEL^, Hap3^INS283^) across 275 tea accessions. Hap1 is present in 66% of CT accessions but absent in cultivated populations, showing a gradual transition from Hap1 to Hap2 and Hap3. **d**, TPM expression levels of *ANS3* across CT, CSA, and CSS, measured in seven accessions per group with three biological replicates each. *ANS3* expression is significantly higher in CT than in cultivated tea plants.

Additionally, we identified shared selection signatures in *UGT91A5* (UDP-glycosyltransferase) and *ANS3* (anthocyanin synthase). The *ANS3* promoter exhibited three SV-derived haplotypes: Hap1 (192 bp insertion), Hap2 (deletion), and Hap3 (283 bp insertion), with distinct population distributions. Hap1 was predominant in CT (66%) but entirely absent in cultivated tea, while CSA accessions were dominated by Hap2 (65%) with moderate Hap3 presence (20%). In contrast, CSS populations displayed Hap3 predominance (71%) with minimal Hap2 retention (4%) (Fig. 4b, c). These results suggest that domestication led to the gradual loss of Hap1, while the CSA-to-CSS transition was characterized by a shift from Hap2 to Hap3. Transcriptomic profiling further revealed significantly higher *ANS3* expression in CT relative to CSA and CSS, with no significant difference between the latter two (Fig. 4d).

The wild relatives (CT) exhibit distinct purple shoots, whereas cultivated tea plants typically display green shoots (Fig. 5a), a phenotypic difference likely governed by anthocyanin metabolism. Previous studies^25^ have reported a strong correlation between anthocyanin content and *ANS3* expression. PCR amplification confirmed the presence of a 192 bp insertion in the *CtANS3* promoter of CT accessions (Fig. 5a). Quantitative Real-Time PCR (qRT-PCR) analysis revealed that *CtANS3* expression was 52-fold higher than in cultivated accessions (Fig. 5b). Consistently, anthocyanin compounds, including delphinidin and cyanidin, were significantly more abundant in CT than in cultivated tea plants (Fig. 5c; Supplementary Fig. 7).

**Fig. 5.**
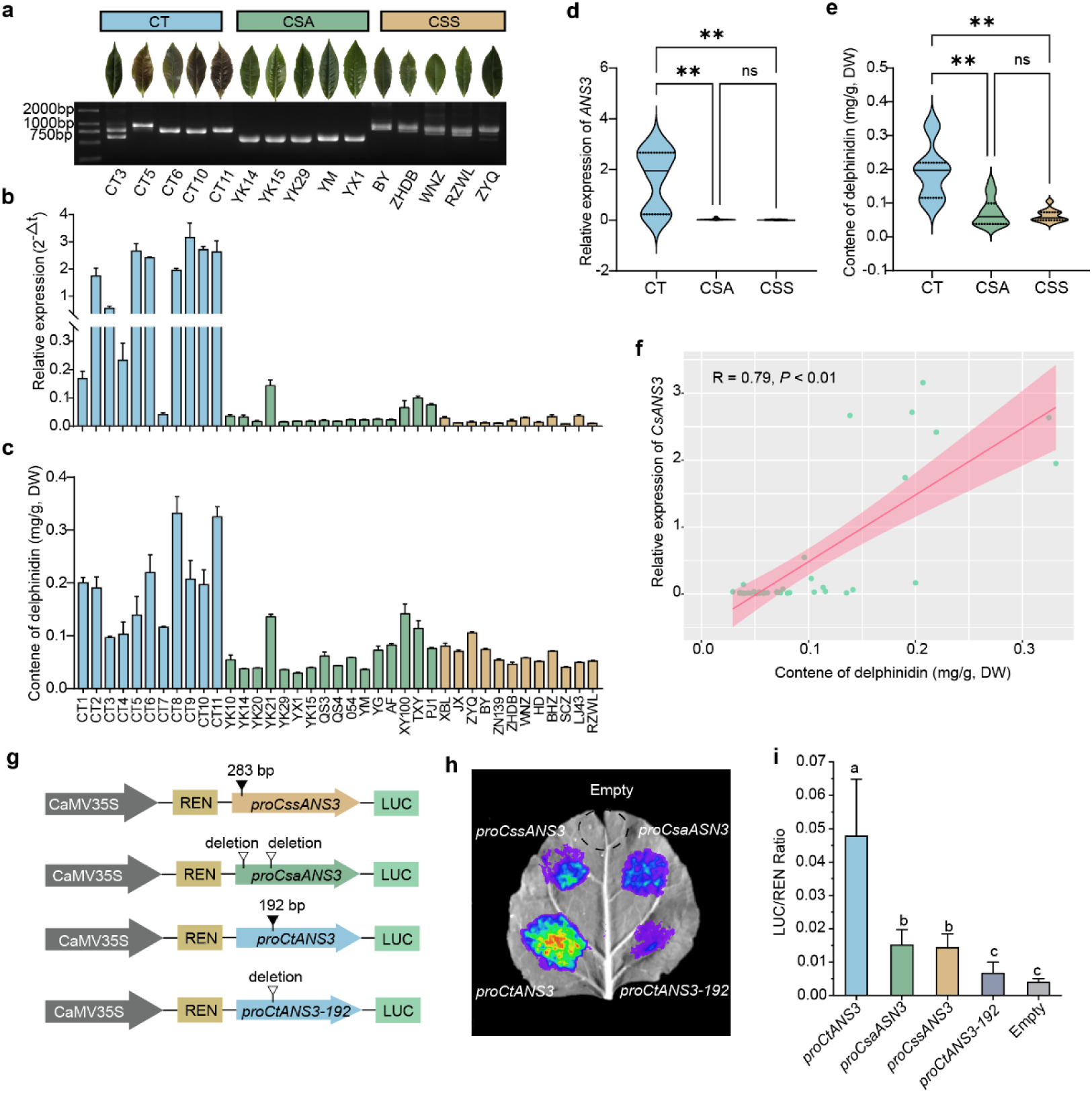
| Effects of SVs in the *ANS3* promoter on gene expression and downstream metabolites. **a**, Genotyping of *ANS3* in 15 accessions using a 2000 bp DNA marker. PCR analysis confirmed the presence of a 192 bp insertion in five CT accessions and a 283 bp insertion in CSS accessions. Experiments were repeated independently three times with consistent results. **b**, qRT-PCR analysis of *ANS3* expression in 39 tea accessions (CT, n = 11; CSA, n = 16; CSS, n = 12), each with three biological replicates. **c**, HPLC quantification of delphinidin in 39 accessions, with DW indicating dry weight. **d, e**, Box plots showing that both *ANS3* expression and delphinidin content are significantly higher in CT accessions, while lowest in CS accessions (\*\**P* < 0.01, one-way ANOVA). **f**, Correlation between *ANS3* expression and delphinidin content across CT, CSA, and CSS accessions. **g**, Schematic of the dual-luciferase assay used to assess *ANS3* promoter activity. **h**, **i**, Luminescence images (**h**) and luciferase (LUC) activity (**i**) showing that *proCtANS3* exhibits significantly higher promoter activity than *proCsaANS3* and *proCssANS3* in transient *N. benthamiana* leaves expression assays. Truncating the 192 bp insertion significantly reduces *proCtANS3*-*192* activity. Relative LUC activity was normalized to REN activity. Eempty represents the empty vector. Data are shown as mean ± SD (n = 4), with significant differences determined using Fisher’s Least Significant Difference (LSD) test.

To investigate the regulatory role of the 192 bp insertion in *CtANS3* expression, we performed promoter activity assays. The *CtANS3* promoter exhibited significantly higher activity than the *CsANS3* promoter from cultivated tea plants. Notably, deletion of the 192 bp fragment substantially reduced promoter activity, confirming its crucial role in driving *ANS3* expression differences between CT and cultivated tea (Fig. 5i). Conversely, CSS accessions harbored a distinct 283 bp insertion (*CssANS3*), but this did not significantly impact *ANS3* expression, anthocyanin accumulation, or promoter activity compared to CSA (Fig. 5b-i). These findings underscore the functional significance of promoter SVs in shaping tea anthocyanin metabolism and domestication-driven phenotypic variation.

### Phenotypic and gene expression characterization of tea plant resistance against *C. gloeosporioides* infection

To assess disease resistance variation among tea accessions, we conducted controlled infection assays using *Colletotrichum gloeosporioides*. Lesion area measurements revealed significantly larger lesions in CT accessions compared to cultivated varieties, whereas no significant difference was observed between CSA and CSS (Fig. 6a-c, Supplementary Fig. 8). Plant resistance to pathogens is largely mediated by resistance (R) genes^26^. We therefore performed a comprehensive analysis of resistance gene analogs (RGAs) across 21 tea plant genomes, categorizing them into seven families based on protein domain architecture (Fig. 6d). Among these, NB-ARC domain-containing RGAs with leucine-rich repeats (LRR) were predominant. Notably, cultivated accessions exhibited a marked enrichment of several RGA classes, including NBS-LRR-TIR, NBS-LRR, CC-NBS-LRR, and RLK-LRR, with NBS-LRR-TIR being 2.3-fold more abundant than in CT. The most prevalent RGA family was RLK-LRR (27.28%), followed by NBS-LRR (25.95%) and CC-NBS-LRR (24.08%), whereas CC-NBS and NBS-TIR were the least represented (3.95% and 1.11%, respectively). Comparative genomic analysis revealed that five gene families, including OG0000028 and OG0000053, exhibited significantly higher copy numbers in cultivated accessions compared to CT, suggesting that R gene expansion may have contributed to enhanced disease resistance in cultivated tea plants.

**Fig. 6.**
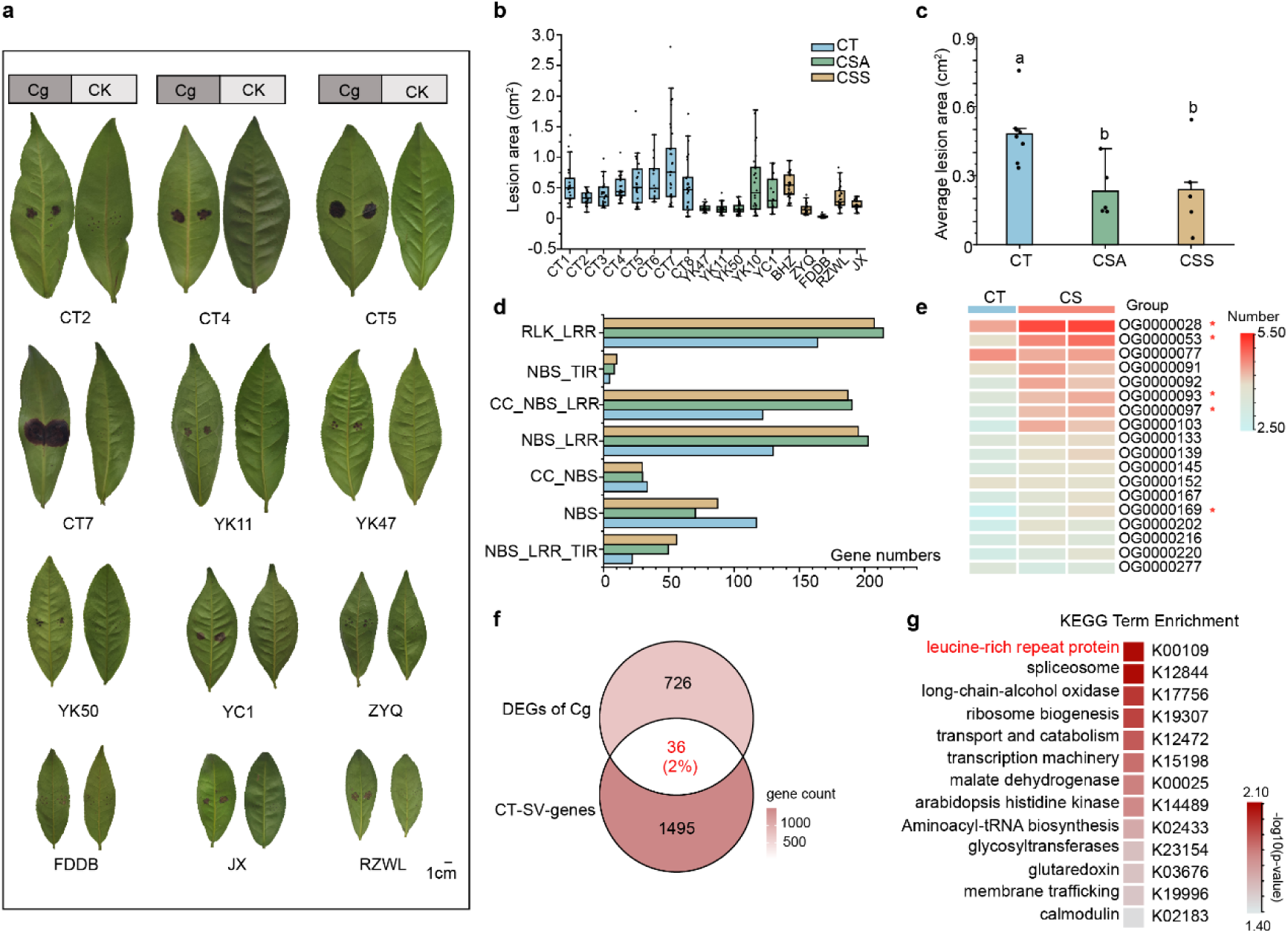
| Phenotypic and transcriptomic characterization of tea plant resistance against *Colletotrichum. gloeosporioides* infection. **a**, Lesion phenotypes in CT, CSA, and CSS accessions following *C. gloeosporioides* (Cg) infection. CT2, CT4, CT5, and CT7 belong to the CT group; YK11, YK47, YK50, and YC1 belong to the CSA group; ZYQ, FDDB, JX, and RZWL belong to the CSS group. CK represents mock infection with water. **b**, Statistical analysis of lesion areas in 18 tea accessions post-infection, with each point representing one replicate (≥15 replicates per accession). **c**, Comparative lesion area analysis among CT, CSA, and CSS accessions. **d**, Statistical classification of R genes across CT, CSA, and CSS genomes, identifying seven RGA types. **e**, Orthogroup-based statistical comparison of gene family sizes in CT, CSA, and CSS genomes. The top 20 gene families showing significant expansion in cultivated tea (*Camellia sinensis*, CS) are shown, with five RGA families marked by red asterisks. **f**, Venn analysis of two gene sets: DEGs of Cg (differentially expressed genes identified from transcriptome analysis of seven accessions post-Cg infection: CT1, CT4, CT7, YK11, YK47, FDDB, and ZYQ) and CT-SV genes (genes in the CT genome with SVs in their promoter regions). **g**. KEGG pathway enrichment analysis of the 36 intersecting genes identified in (**f**).

Structural variant (SV) analysis of R genes in the reference genome further revealed extensive genomic variation within this gene class. Of the 739 identified R genes, 398 (53.85%) harbored SVs within their mRNA regions, averaging five SV events per gene locus (Supplementary Fig. 9, Supplementary Table 10). Additionally, SVs were detected in the promoter regions of 326 R genes (44.11% of the total RGA repertoire), indicating frequent structural modifications in regulatory elements (Supplementary Table 10).

### A 159 bp regulatory insertion upregulates *CtLRR1* and impairs pathogen defense

To elucidate the molecular mechanisms underlying disease susceptibility in wild tea relatives, we performed transcriptome sequencing on seven pathogen-infected accessions. Comparative analysis of differentially expressed genes (DEGs) between cultivated and CT accessions identified 129 overlapping DEGs in cultivated accessions and 674 in CT accessions, which were combined into a non-redundant set of pathogen-responsive genes (Supplementary Fig. 10). To assess whether structural variants (SVs) contribute to disease susceptibility in CT accessions, we conducted an integrative analysis comparing SV-associated genes (CT-SV) with this pathogen-responsive gene set. This analysis revealed 36 differentially expressed genes harboring SVs (Fig. 6f). Gene ontology (GO) enrichment indicated a significant association with leucine-rich repeat (LRR) domain-containing proteins (Fig. 6g).

Consequently, we identified a 159 bp specific insertion in the promoter region of Among these, we identified a 159 bp insertion in the promoter region of *CtLRR1* specific to CT accessions (Fig. 7a). PCR validation confirmed that this insertion is heterozygous in CT but a homozygous deletion in cultivated accessions (Fig. 7b). Following pathogen infection, *CtLRR1* expression was significantly upregulated in susceptible accessions compared to resistant ones (Fig. 7c). Luciferase reporter assays further supported this finding, showing that *CtLRR1* promoter activity was significantly higher than that of *CsaLRR1* and *CssLRR1* (Fig. 7e).

**Fig. 7.**
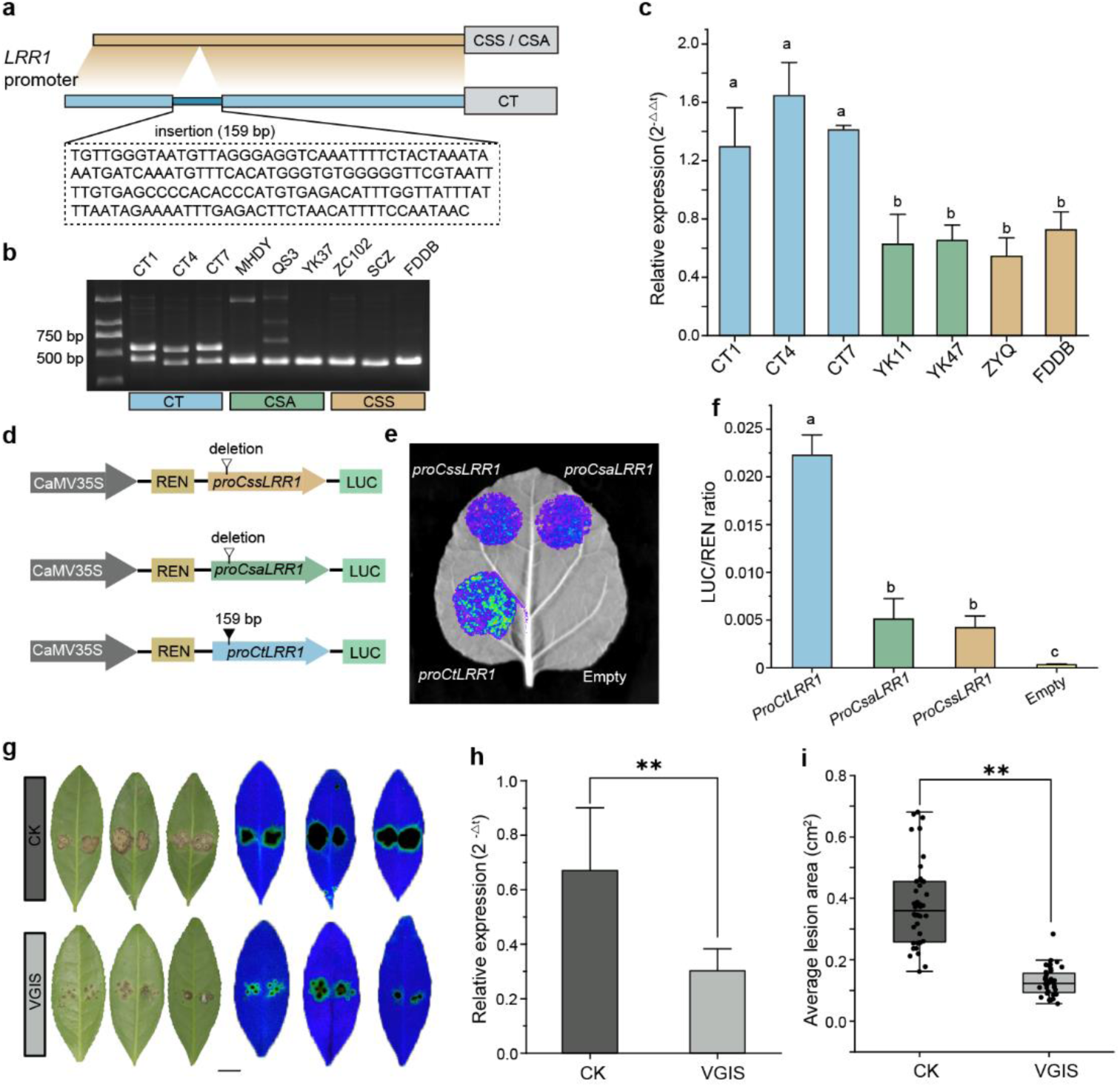
| Functional analysis of the 159 bp *CtLRR1* promoter insertion in pathogen defense. **a**, Schematic representation of the SV in the *LRR1* promoter. **b**, PCR verification of *LRR1* promoter SVs in seven tea accessions. **c**, qRT-PCR analysis of *LRR1* expression in seven tea accessions. **d**. Diagram illustrating the dual-luciferase assay used to assess *LRR1* promoter activity. **e**, **f**, Luminescence imaging (**e**) and luciferase (LUC) activity quantification (**f**) show that transient expression of *CtLRR1* in *N. benthamiana* leaves results in higher promoter activity than *CsaLRR1* and *CssLRR1*. Relative LUC activity was normalized to REN activity. Empty represents the empty vector control. Data are presented as mean ± SD (n = 4 biological replicates). **g**, CK represents tea leaves infected with *Agrobacterium* carrying empty PTRV1 and PTRV2 vectors, while VIGS represents tea leaves infected with *Agrobacterium* carrying empty PTRV1 and *CsLRR1*-PTRV2. The right panel shows chlorophyll fluorescence imaging of infected leaves. **h**, *CsANS3* expression levels in control and silenced samples. **i**, Lesion area measurements following *C. gloeosporioides* (Cg) infection in control and VIGS-silenced groups (n = 40 per group). \*\**P* < 0.01 (one-way ANOVA).

To investigate the functional role of *LRR1*, we employed virus-induced gene silencing (VIGS) to suppress its expression in tea plants. Compared to pathogen-infected controls, silenced plants exhibited significantly reduced leaf damage (Fig. 7f-i). These results indicate that the 159 bp insertion in the *CtLRR1* promoter enhances promoter activity and transcriptional levels but impairs pathogen defense, suggesting a negative regulatory role in CT accessions’ resistance to *Colletotrichum gloeosporioides*.

## Discussion

Limited genomic data on wild relatives and *Camellia sinensis* var. *assamica* (CSA) has hindered comprehensive research and the effective utilization of tea plant genetic resources. To address this, we assembled a new wild relative genome and five haplotype genomes of cultivated tea plants (three CSA and two CSS) using PacBio HiFi sequencing. Compared to existing references, our genome assemblies exhibit superior contiguity (contig N50), completeness (BUSCO scores), and structural integrity (LAI scores). Using these high-quality resources, we constructed a comprehensive pangenome from 21 tea plant genomes, selecting previously published assemblies with robust quality and broad geographical representation. Through extensive structural variant (SV) analysis, population-level SV studies, and transcriptome sequencing, we elucidated how SVs influence gene expression during tea domestication, shaping morphological diversity across accessions.

The divergence time between wild and cultivated tea plants has been widely debated. Previous studies estimated a divergence of ∼24 MYA between DASZ and cultivated tea plants^27^; however, our analysis, incorporating a broader genomic dataset with higher-quality assemblies, refined this estimate to ∼5.84 MYA (Fig. 1a). Given *C. sinensis*’s high heterozygosity and genetic diversity, an accurate pangenome must encompass representatives with significant genetic divergence (*e.g*., CSA and its wild relatives). Our pangenome reaches saturation in both pangenes and core genes, demonstrating greater completeness than previously reported tea plant pangenomes ^20,21^. Comparative gene family analysis revealed that CSA and CSS accessions underwent expansions in 153 and 223 gene families, respectively, with significant enrichment in defense-related pathways, particularly those involved in disease and insect resistance. This expansion contrasts with the loss of defense-related genes observed during grape domestication^28^, suggesting that tea domestication favored stress resistance.

SVs are key drivers of plant phenotypic diversity and genome evolution^29–32^. Analyzing 21 tea plant genomes, we identified 522,428 non-redundant SVs, predominantly large insertions and deletions (>50 bp, 75.36%), followed by translocations (25.07%) and inversions (1.37%). Interestingly, unlike rice and broomcorn millet, where wild species exhibit higher SV frequencies^33,34^, ur data reveal lower SV frequencies in wild relatives than in cultivated accessions. This discrepancy may stem from alignment biases due to the elevated inversion and translocation frequencies in wild populations. Moreover, insertions and deletions were more prevalent in promoter regions than in coding or intronic sequences, consistent with patterns observed in kiwifruit^35^. Transcriptome analysis further linked promoter-region SVs to differential gene expression in flavonoid, amino acid, and terpenoid metabolism, likely contributing to flavor diversity among tea accessions. Comparable SV patterns have been reported in other crops; for example, wheat pangenome analysis identified a large insertion/deletion block on chromosome 1RS with reduced *pSc200* tandem repeats in cultivars, a region widely utilized in breeding programs^36^. Such parallels underscore the importance of insertion/deletion variations in crop improvement. To support future investigations, we cataloged SVs affecting gene promoter regions within our pangenome (Supplementary Table 13), enabling targeted analyses of gene families or key regulatory loci linked to critical quality traits.

In diploid plants, SV-induced allele loss results in hemizygous genes, a largely unexplored phenomenon in tea^37–39^. By assembling 10 haplotype genomes, we identified hemizygous genes arising from SVs. Notably, CSA accessions exhibited slightly higher hemizygosity (3.96%) than CSS accessions (3.39%), with these genes predominantly involved in primary cell wall biosynthesis and ovule development. This pattern mirrors findings in grape^28^, where cultivated accessions display reduced hemizygous gene frequencies.

To harness the genetic diversity captured in our pangenome, we integrated SV data from 275 tea accessions into a comprehensive variation map, constructing a graph-based reference genome encompassing both wild and cultivated tea plants. Such graph-based frameworks have proven invaluable for SV-based association studies in other crops, enabling the identification of key agronomic traits^40–42^. However, due to the absence of extensive phenotypic datasets, we relied on selection sweep analyses to delineate domestication-associated genomic intervals. Comparative scans between CT and CSS populations revealed selective pressures on resistance– and adaptation-related genes, including *OFP*, *COR*, and *RPV*, aligning with recent findings in tea^16^. Notably, during the transition from CSA to CSS, selection targeted key flavor-related genes, including *HCT*, *CHI*, *LAR*, and *TPS*. Among these, the anthocyanin biosynthesis gene *ANS3* underwent convergent selection. Integrating genotype data, expression profiles, and functional validation, we demonstrated that a CT-specific 192 bp promoter insertion significantly enhanced *CtANS3* transcriptional activity, leading to elevated anthocyanin accumulation. This regulatory mechanism, wherein promoter-region SVs drive gene expression, has also been observed in apple^14^ and cabbage^43^, suggesting a conserved role in plant domestication.

Research on disease resistance in tea plants remains a key focus in the genetic improvement of domesticated cultivars. Consistent with findings in rice^33^ and chickpea^12^, our study revealed a greater abundance of resistance gene analogs (RGAs) in cultivated tea accessions than in their wild relatives, with a notable 2.3-fold expansion in NBS-LRR-TIR class genes. This pattern parallels R-gene enrichment during domestication in apple^26^, suggesting that artificial selection broadly drives the expansion of crop resistance genes. Recent chickpea pangenome studies^12^ have shown that SVs are prevalent in resistance (R) genes, with an average of three SV events per gene. Similarly, we observed frequent SVs within R gene bodies and identified that over 44% of R gene promoter regions contained SVs. By integrating pathogen transcriptomic data from different tea accessions, we pinpointed a heterozygous site in CT accessions caused by a 159 bp insertion in the *CtLRR1* promoter. Functional validation demonstrated that while this insertion significantly upregulated *CtLRR1* transcription in CT accessions, it paradoxically reduced resistance to *Colletotrichum gloeosporioides*. In contrast, cultivated tea plants lacking this insertion exhibited enhanced resistance. Similar antagonistic regulatory effects have been reported in *Arabidopsis*^44^, underscoring the complexity of disease-resistance gene networks. These findings advance our understanding of disease resistance mechanisms in tea domestication and highlight the regulatory roles of SVs in crop evolution.

In summary, we present six chromosome-level reference genomes and construct a graph-based whole-genome SV map for tea plants. Our comprehensive SV analysis reveals differential gene expression patterns linked to key agronomic traits, including leaf coloration, flavor compound biosynthesis, and disease resistance. More importantly, by integrating pangenomic and transcriptomic data, we systematically identify SVs across tea populations and demonstrate their bidirectional regulation of multiple gene expression levels. Notably, many of these expression-modulating SVs have undergone strong selection pressure during domestication. The genomic resources and pangenome variation map provided in this study offer valuable tools for future crop improvement, paving the way for accelerated advancements in tea plant breeding.

## Methods

### Genome sequencing and assembly

We sequenced and assembled the genomes of five tea accessions representing two *Camellia sinensis* varieties: three *C. sinensis* var. *assamica* (MHDY, QS3, and YK37) from Xishuangbanna Prefecture, Yunnan, and two *C. sinensis* var. *sinensis* (HJY and ZC102) from Anhui, China. Genomic DNA was extracted from fully expanded mature leaves and used to construct PacBio SMRT libraries with the SMRTbell Express Template Prep Kit 2.0. Libraries were sequenced on the PacBio Sequel II platform using the Circular Consensus Sequencing (CCS) model^45^, with six cells per sample, yielding >16K reads per cell and an average coverage depth of 38.13×. In addition to these five accessions, the genome of one wild relative (DLC) was previously sequenced using the Continuous Long-Read (CLR) technology, but the data has not been publicly released. Hi-C libraries were prepared using standard protocols and sequenced on the DNBseq platform. Raw reads were filtered with SOAPnuke to remove low-quality data. Hifiasm^46^ was employed to generate haplotype-resolved genome assemblies by integrating HiFi and paired-end Hi-C data. This process produced two haplotype-specific contig sets and a primary assembly, which was scaffolded into chromosome-level reference sequences using Hi-C data^47^. Haplotype-specific assemblies were aligned to the reference using RagTag^48^ to obtain chromosome-scale haplotype assemblies (Supplementary Note).

Whole-genome resequencing was conducted on 275 tea accessions, including 94 wild relatives, 97 *C. sinensis* var. *assamica* (CSA), and 84 *C. sinensis* var. *sinensis* (CSS) accessions. Of these, 32 accessions were from previous studies, while 243 were newly sequenced. Detailed accession information is provided in Supplementary Table 11.

### Repeat sequence annotation

Repetitive sequences were annotated using both homology-based and *de novo*^49^ approaches. The homology-based method utilized the RepBase^50^ database, applying RepeatMasker and RepeatProteinMask to identify and classify known repetitive elements. For *de novo* prediction, we constructed a repeat library using RepeatModeler^51^ and LTRharvest, followed by annotation with RepeatMasker. Tandem repeats were identified with Tandem Repeats Finder.

### Gene prediction and functional annotation

Gene structures were predicted by integrating homology-based and transcriptome-assisted annotation. Homology-based prediction involved aligning protein-coding sequences from related species to the target genome using GeMoMa^52^. Transcriptome-assisted annotation was conducted using RNA-seq and ISO-seq data. RNA-seq reads were aligned with HISAT2^53^ and assembled with StringTie^54^, while ISO-seq reads were processed with GMAP^55^ and PASA^56^ assembly. GeMoMa was used to generate a final non-redundant gene set, which was evaluated via BUSCO^57^ for completeness.

Functional annotation was performed by aligning gene sequences against SwissProt^58^, TrEMBL^58^, NR, and KOG using Diamond software, followed by biological pathway annotation using KEGG^59^ and gene ontology annotation using GO^60^. Detailed genome prediction methods in the supplement note.

### Phylogenetic tree construction and divergence time estimation

The *Actinidia chinensis* cv. ‘Hong Yang’ (v3) genome (https://kiwifruitgenome.org/) was used as an outgroup for phylogenetic inference. Single-copy orthologs were identified using OrthoFinder (v2.5.5)^61^ with the “-S blast” parameter. Protein sequences of single-copy genes were aligned with MAFFT (v7.525)^62^, and codon alignments were generated using PAL2NAL (v14)^63^ with default parameters. Conserved alignment blocks were extracted using Gblocks (v0.91b) with parameters set to ‘-t=c’ for codon-based selection and ‘-b5=h’ for allowed gap positions. The resulting filtered alignments were then concatenated into a supermatrix for downstream analyses. Phylogenetic reconstruction was performed using IQ-TREE (v2.3.3)^64^ with parameters ‘-st CODON –bb 1000 –m GY+F+G4’. Divergence time estimation was conducted using the MCMCtree program implemented in PAML (v4.9j)^65^. Two fossil calibration points were employed: the divergence between *Actinidia chinensis* and *Camellia sinensis* (82.8-106 Ma), and the split between *Camellia tailensis* and *Camellia sinensis* (4.8-7.8 Ma) from Time Tree (https://timetree.org/home) database.

### Analysis of gene gain and loss

According to the clustering results, we identified 45,780 non-redundant orthogroups, which were subsequently classified based on their distribution across samples: gene families of core (present in all 21 samples), soft-core (19-20 samples), dispensable (2-18 samples), and private (unique to a single sample). Pan-genome saturation curves were generated using PanGP (v1.0.1) (https://pangp.zhaopage.com/) to visualize the gene clustering results. After calculating the number of genes per genome in each gene family, we divided the samples into three groups (CT, CSA, and CSS) for comparative analysis. Statistical significance was assessed using the Mann-Whitney-Wilcoxon test with a significance threshold of *P* < 0.05^28^. Genes showing significantly higher copy numbers in a group were classified as gains, while those with significantly lower copy numbers were classified as losses. Gene Ontology (GO) enrichment analysis was performed using the topGO package in R.

### Identification of disease-resistance genes

Based on established methodology^66,67^, we identified and classified disease resistance genes in the tea plant genome, focusing on two major types: NBS-LRR (nucleotide-binding site with leucine-rich repeat) and RLK-LRR (pattern-recognition receptor) genes. Using HMMER with default parameters, we searched the tea plant proteome against the Hidden Markov Model (HMM) of the NB-ARC domain (PF00931). The identified NBS-encoding genes were further classified by searching against TIR (PF01582) and LRR domain HMMs from the PFAM database. Additionally, CC domains within these NBS-encoding proteins were identified using ncoils under default settings. For RLK-LRR gene identification, we first searched the tea plant proteome using the kinase HMM profile (PF00069) from PFAM database, followed by scanning the resulting candidates against the LRR HMM profile (PF00560).

### Detection of genomic structural variants

The whole genome alignment between twenty tea plant genomes and the ZC102 reference was performed using minimap2 (v2.17)^68^ with parameters ‘-ax asm5 –eqx’, followed by BAM file sorting using samtools. Structural variants (SVs) were identified using SyRI (v1.6)^69^, including insertions (>50 bp), deletions (>50 bp), inversions, and translocations. We filtered out SVs containing ‘N’ sequences, those with ambitious alignment margins, and variants showing poor synteny alignment at breakpoints. SURVIVOR (v.1.0.7)^70^ was employed to merge structural variants (SVs) shorter than 50 kb across all analyzed genomes. The consolidation was performed using the parameter set ‘1000 2 1 0 0 50’, which specified a maximum breakpoint distance of 1,000 bp for merging SVs from the input VCF files. The distribution of identified SVs was visualized using Circos (v0.69-8)^71^. Identification of hemizygous genes was performed by detecting SVs between haplotype 1 and haplotype 2 using minimap2 followed by SyRI analysis. The genes surrounded by heterozygous insertion/deletion variations were identified as hemizygous candidates^28,72^, and their quantification was performed using bedtools (v2.30.0)^73^.

### SV-affected genes and associated expression analysis

SVs overlapping with CDSs, introns, and promoter regions (defined as 2,000 bp upstream of each gene) were identified using bedtools intersect (’-wa –wb’). After removing redundancies, genes associated with SVs were classified as SV-affected. RNA sequencing was conducted on 21 tea accessions categorized into CT, CSA, and CSS groups. Transcriptomic data were used to determine gene expression levels, and SV-affected genes were compared across groups. Genes exhibiting a >2-fold expression change with TPM > 1 in >60% of samples were selected for heatmap visualization and enrichment analysis. Expression patterns were visualized using TBtools^74^ and KEGG pathway enrichment analysis was performed using clusterProfiler^75^.

### Graph-based tea plant genome construction and SV genotyping

We generated a graph-based representation of the tea plant genome by integrating the linear reference sequence with large-scale genomic variants through the implementation of ‘vg construct’ pipeline^76^. The merged VCF files were compressed and indexed using bcftools, followed by the implementation of ‘vg giraffe’ through EVG^77^ installation. A genome graph was constructed and indexed based on the ZC102 reference genome using ‘vg autoindex’ and ‘vg snarls’. Using ‘vg giraffe’, we mapped short paired-end reads derived from 275 tea accessions against the indexed graph genome, resulting in the generation of GAM format alignment files. The graph alignment results were processed using ‘vg pack’ to generate compressed coverage information, followed by structural variant (SV) genotyping of all 275 tea accessions through the implementation of ‘vg call’ using default parameters.

### Selective sweep analysis

Selective sweeps were identified by merging all gVCF files using bcftools ‘merge’^78^. Nucleotide diversity (π) was calculated separately for CT, CSA, and CSS populations using vcftools^79^, and pairwise *F*_st_ comparisons were conducted with a sliding window approach (parameters: ‘--fst-window-size 100000 –-fst-window-step 10000’). The top 5% of *F*_st_ regions were designated as putative selective sweeps.

### Quantification of Anthocyanin Content

Anthocyanin extraction and quantification were performed as previously described^80^. Lyophilized tea leaves (50 mg) were ground into powder and suspended in 1 mL of 1% HCl/methanol (v/v). After sonication at 90% power for 30 minutes, the mixture was centrifuged at 5,000 rpm for 5 minutes, and the supernatant was collected. The residue was extracted twice with 500 μL of 1% HCl/methanol, and all three supernatants were combined. A 1:1:1 (v/v/v) mixture of the combined extract, ultrapure water, and chloroform was vortexed for 2 minutes, centrifuged at 5,000 rpm for 5 minutes, and the upper phase was collected. This phase was mixed with ultrapure water and ethyl acetate (1:1:1, v/v/v), vortexed for 2 minutes, centrifuged at 5,000 rpm for 5 minutes, and the lower phase containing anthocyanins was collected. To obtain anthocyanin monomers, 2 mL of the extract was transferred to a 15 mL centrifuge tube, combined with 200 μL of concentrated HCl, loosely capped, and incubated in a 100 °C water bath for 1 hour. The extract was cooled on ice for 10 minutes, filtered through a 0.22 μm membrane, and analyzed by UPLC.

### UPLC analysis

The anthocyanins in tea samples were extracted and analyzed following the method of Mei et al.^81^. Ultra-performance liquid chromatography (UPLC) was performed using a Waters ACQUITY UPLC system (Waters, USA) equipped with a Phenomenex Kinetex C18 column (2.6 μm, 100 mm × 4.6 mm i.d.). The mobile phase consisted of 0.5% formic acid (v/v) in water (A) and acetonitrile (B), with a gradient elution program as follows: 8–20% B over 4 minutes, held at 20% B for 2.5 minutes, 20–40% B over 5.5 minutes, held at 40% B for 1 minute, 40–90% B over 2 minutes, and re-equilibrated to 10% B over 5 minutes, totaling a 20-minute run time. The flow rate was maintained at 0.8 mL/min. Detection was performed at 530 nm, with the column and autosampler temperatures set at 25 °C and 4 °C, respectively. Anthocyanin monomers were identified and quantified using standard curves generated from cyanidin, delphinidin, and pelargonidin. All measurements were performed in biological triplicate, with means and standard deviations calculated accordingly.

### DNA extraction and PCR verification

Genomic DNA was extracted from 100 mg of tea leaves ground in liquid nitrogen. The lysate was prepared using 500 μL of extraction buffer and 10 μL β-mercaptoethanol, incubated at 65 °C for 30 minutes, followed by the addition of 600 μL chloroform/isoamyl alcohol (24:1 v/v) and centrifugation at 12,000 rpm for 10 minutes. The supernatant was mixed with an equal volume of isopropanol, centrifuged at 12,000 rpm for 5 minutes, and the resulting pellet was washed twice with 70% ethanol before resuspension in 100 μL of sterile water. Primers were designed based on syntenic regions across 21 genomes, and gene fragments were validated via PCR and agarose gel electrophoresis. All primer sequences are listed in Supplementary Table 12, and sequencing data are provided in the Supplementary Note.

### Quantitative real-time PCR

Total RNA was extracted from young tea leaves using a Plant RNA Kit (RC411-01, Vazyme, China), and first-strand cDNA was synthesized with a HiScript III RT SuperMix for qPCR Kit (R323-01, Vazyme, China). qPCR was conducted using Taq Pro Universal SYBR qPCR Master Mix (Q712-02, Vazyme, China) on a BioRad CFX96 Real-Time PCR Detection System (Bio-Rad, USA). The thermal cycling conditions were 95 °C for 3 minutes, followed by 38 cycles of 95 °C for 10 s and 57 °C for 30 s, then 65 °C for 5 s and 95 °C for 5 s. *CsGAPDH* was used as an internal reference, and relative gene expression was calculated using the 2^−ΔCT^ method. For samples treated with Colletotrichum (with CK controls), the 2^−ΔΔCT^ method was used.

### Dual-luciferase reporter assay

The promoter sequences of *LRR1* and *ANS3* were cloned from different tea cultivars, and a 192 bp insertion in the *CtANS3* promoter was removed using fusion PCR. In total, seven promoters were obtained: *proCssLRR1*, *proCsaLRR1*, *proCtLRR1*, *proCssANS3*, *proCsaANS3*, *proCtANS3*, and *proCtANS3-192*. These promoter sequences were inserted into the pGreenⅡ-0800-LUC vector via homologous recombination (restriction sites: *Kpn Ⅰ – Hind Ⅲ*), with the empty pGreenⅡ-0800-LUC vector serving as a control. The recombinant constructs were transformed into *Agrobacterium tumefaciens* strain EHA105 (pSoup) (Weidi, China), and positive transformants were selected on LB medium containing 50 mg/L kanamycin. The verified *Agrobacterium* cultures were grown, collected by centrifugation at 5,000 rpm for 10 minutes, and resuspended in infiltration buffer (10 mM MES, 10 mM MgCl₂, and 200 μM acetosyringone) to an OD₆₀₀ of 0.4. The bacterial suspensions (100 μL per infiltration site) were introduced into the leaves of six-week-old *Nicotiana benthamiana* plants using a needleless syringe.

Following infiltration, the plants were kept in darkness for 48 hours. Afterward, 100 mM potassium D-luciferin (Biolai, China) was applied to the abaxial leaf surface, and luminescence signals were captured using the NightShade LB 9851 *in vivo* imaging system (Berthold, Germany). Relative luminescence activity was quantified using the Promega E1500 Luciferase Assay System (Promega, USA), calculated as the *Luc/Ren* ratio.

### Pathogen Infection Transcriptome Analysis

Tea plant cultivars with uniform growth were collected from the germplasm repository at Yunnan Academy of Agricultural Sciences. The samples included eight CT accessions (CT1–8), five CSA cultivars (YK47, YK11, YK50, YK10, YC1), and five CSS cultivars (BHZ, ZYQ, FDDB, RZWL, JX). Leaves from these cultivars were inoculated with *Colletotrichum gloeosporioides* (Cg) mycelial plugs, while water-treated leaves served as controls (CK). Each treatment was performed in biological duplicate per leaf. Inoculated samples were sealed with preservative film and incubated in a growth chamber at 25 °C under a 16-hour light/8-hour dark cycle with relative humidity exceeding 80%. Disease progression was monitored daily, and lesions became distinct by day 6 post-inoculation. Leaves were then photographed, and lesion areas were quantified using ImageJ software.

For transcriptome sequencing, seven accessions (CT1, CT4, CT7, YK11, YK47, FDDB, ZYQ) were selected. Gene expression levels were quantified using the HISAT2 and StringTie pipeline with default parameters. Differentially expressed genes (DEGs) were identified using DESeq2, with genes showing fold changes >2 and *P* < 0.05 retained for downstream analysis.

### Virus-Induced Gene Silencing (VIGS)

VIGS was performed following established protocols with modifications optimized for tea plants^82–84^. One-year-old tea seedlings were infiltrated with *Agrobacterium* suspensions carrying either pTRV1/pTRV2 or pTRV1/pTRV2-*CsLRR1* via the leaf injection method. Infiltrated plants were incubated at 25 °C for 14 days, after which *CsLRR1* transcript levels were assessed by qRT-PCR. Following gene silencing confirmation, the treated leaves were inoculated with *C. gloeosporioides* mycelial plugs. After 6 days, lesion areas were quantified using ImageJ, and disease symptoms were examined via chlorophyll fluorescence imaging.

## Data availability

All data generated in this study have been deposited in the National Genomics Data Center (NGDC) (https://bigd.big.ac.cn/bioproject). Genomic and transcriptome data are available under BioProject accession number PRJCA036376. Resequencing data are deposited under BioProject accession number PRJCA036909 and will be publicly accessible upon publication.

## Acknowledgments

This work was supported by the National Natural Science Foundation of China (grant Nns. U20A2045, 32472791, 32260790, 32202542 and 32472791), the Project of Science and Technology of Yunnan Province (grant no. 202102AE090038) and the Base of Introducing Talents for Tea Plant Biology and Quality Chemistry (D20026).

