## Supplementary Figures for "Pangenome analyses of tea plant reveal structural variations driven gene expression alterations and agronomic trait diversification"

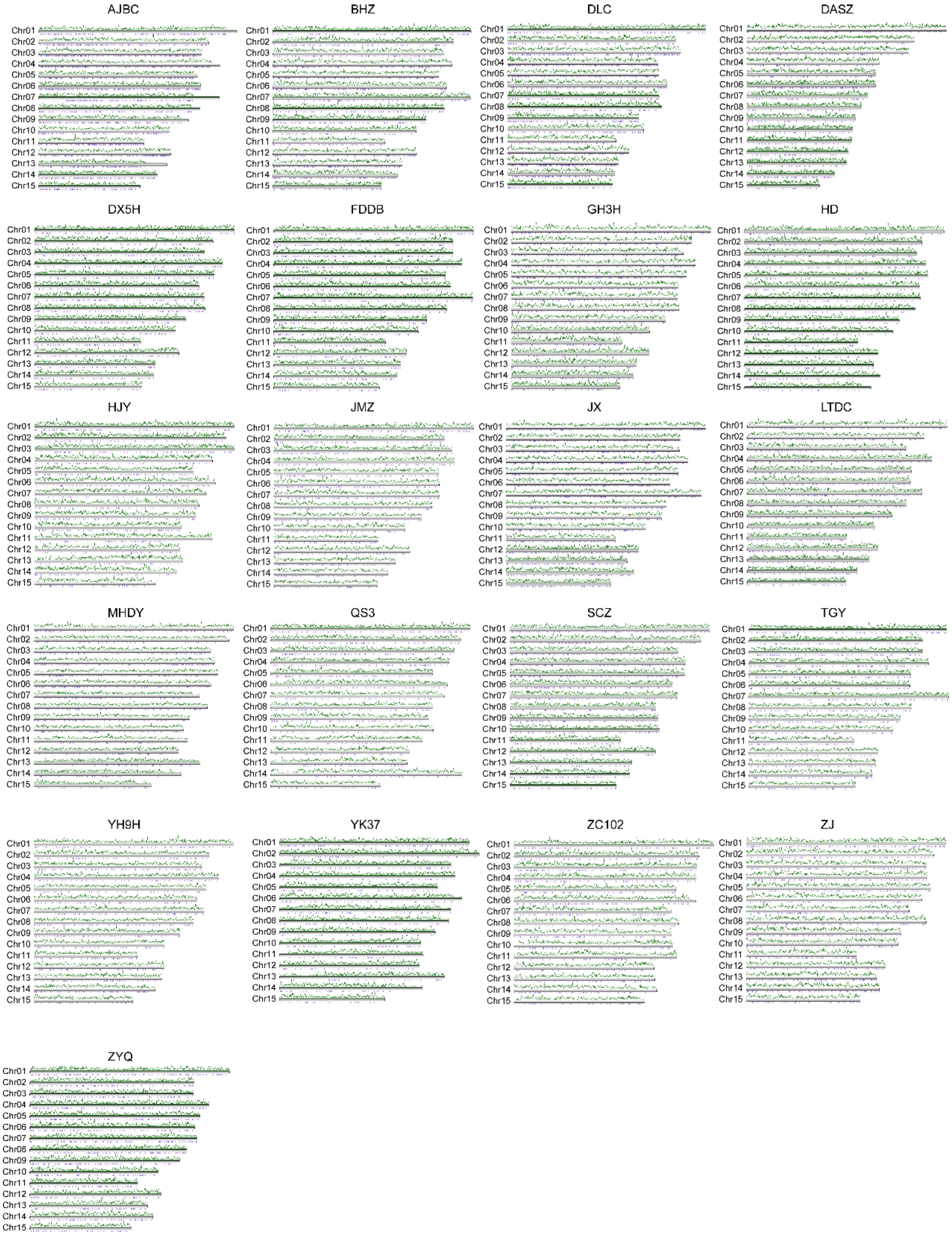


**Supplementary Fig. 1. The distribution of LTRs on 15 chromosomes across 21 genomes.** The green broken line represents the density of LTRs with a window size of 500 Kb, and the purple vertical lines represent the genes with LTR insertions.

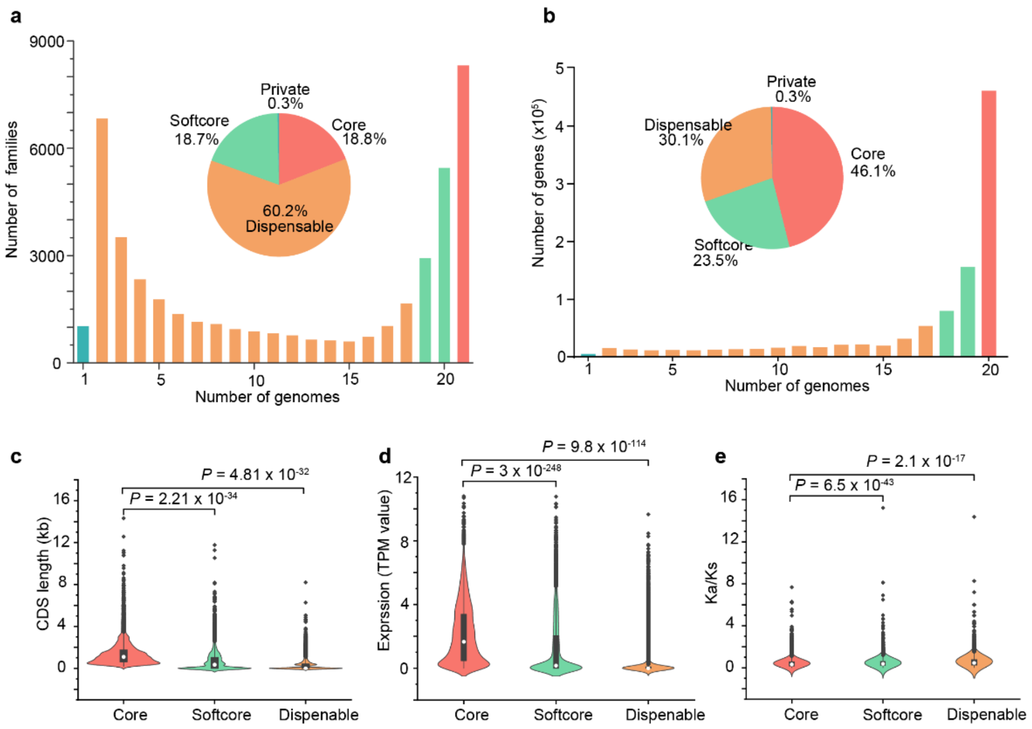


**Supplementary Fig. 2. Pangenome analysis at the gene space.** a) The proportions of pangene families in the core, softcore, dispensable, and private categories. b) The proportions of the number of genes in pangene families within the core, softcore, dispensable, and private categories. c-e) Comparison of CDS lengths, TPM values, and Ka/Ks ratios of core, softcore, and dispensable genes.

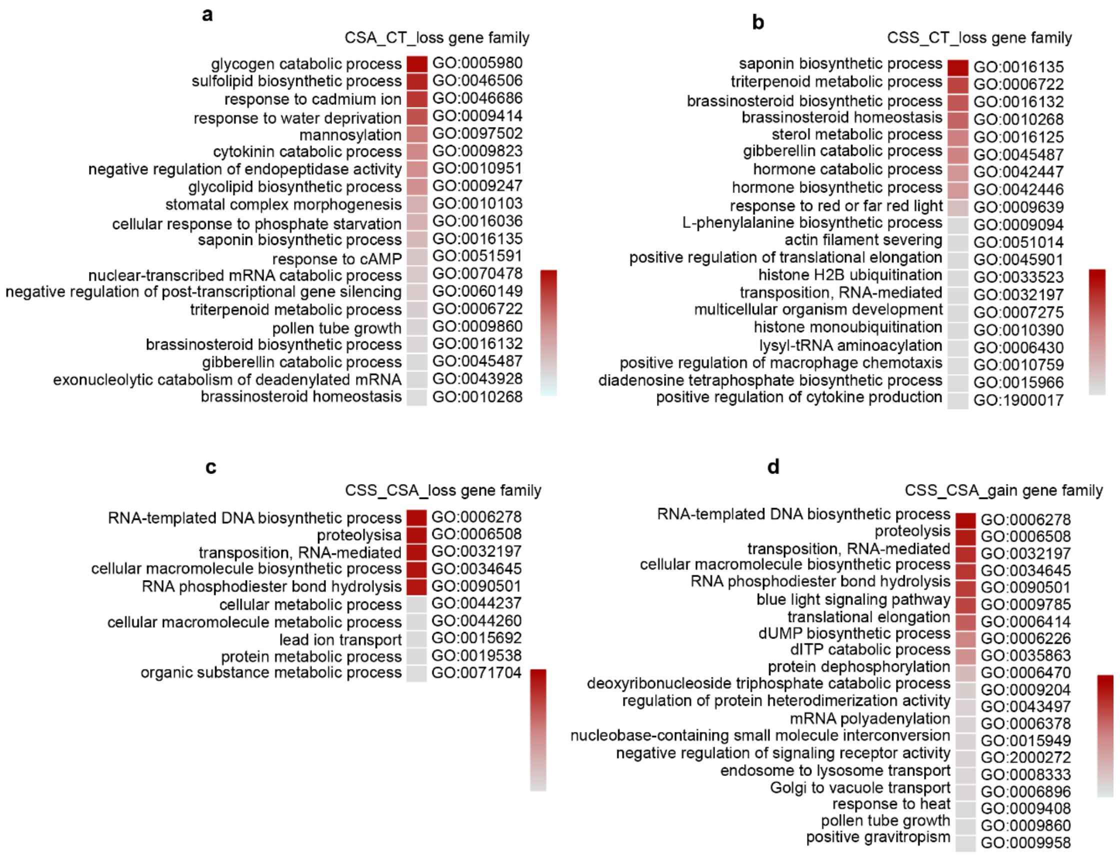


**Supplementary Fig. 3. The gene gain and loss during the domestication of tea plants.** a) GO enrichment results for genes lost by the CSA population compared to CT. b) GO enrichment results for genes lost by the CSS population compared to CT. c) GO enrichment results for genes lost by the CSS population compared to CSA. d) GO enrichment results for genes gained by the CSS population compared to CSA.

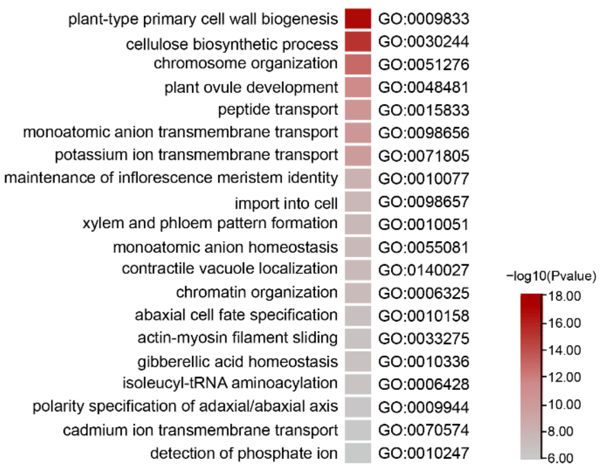


**Supplementary Fig. 4. GO enrichment of hemizygous genes in 10 haplotype genomes.**

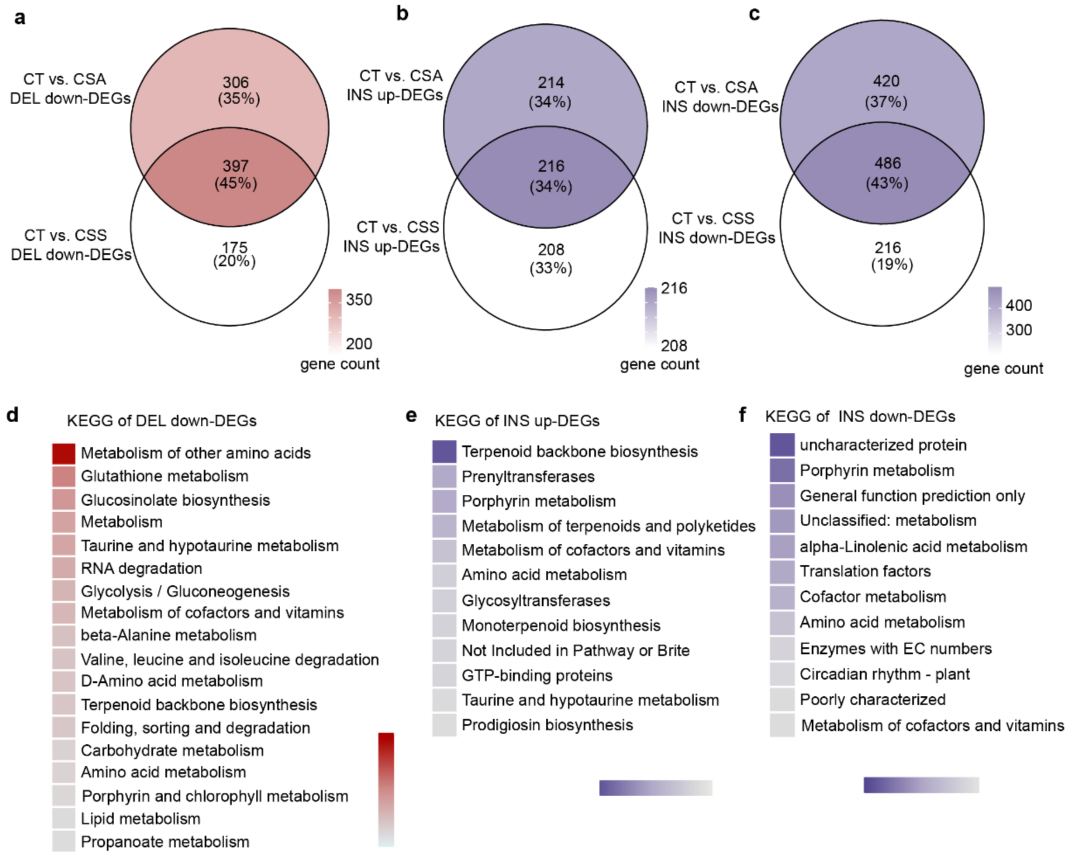


**Supplementary Fig. 5. Insertions and deletions in the promoter region diver differential gene expression among tea plant populations.** a) Genes with deletion mutations in the promoter region that were downregulated in CSA and CSS compared to CT. b) Genes with insertion mutations in the promoter region that were upregulated in CSA and CSS compared to CT. d) Genes with insertion mutations in the promoter region that were downregulated in CSA and CSS compared to CT. d-f) KEGG enrichment results for the overlapping genes corresponding to a-c.

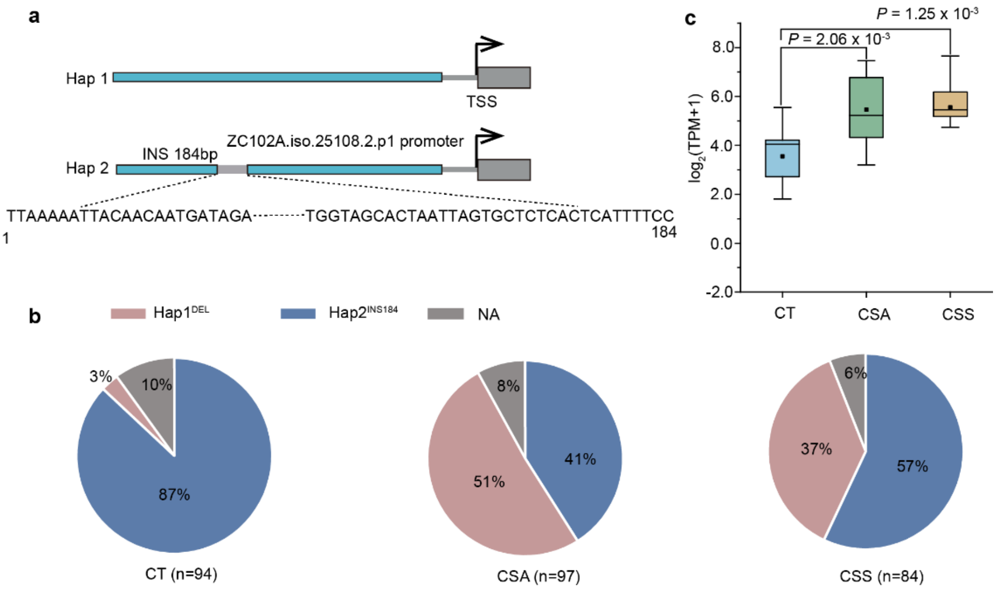


**Supplementary Fig. 6. Haplotype variation in the *LRR1* promoter region and its distribution across tea plant populations.** a) Schematic diagram of the 184bp insertion in the *LRR1* gene promoter region. b) Distribution of hap1 and hap2 across CT, CSA, and CSS populations, showing the gradual decrease of hap2 during domestication from CT to CSA and CSS. c) Expression levels (TPM) of *LRR1* gene in CT, CSA, and CSS populations.

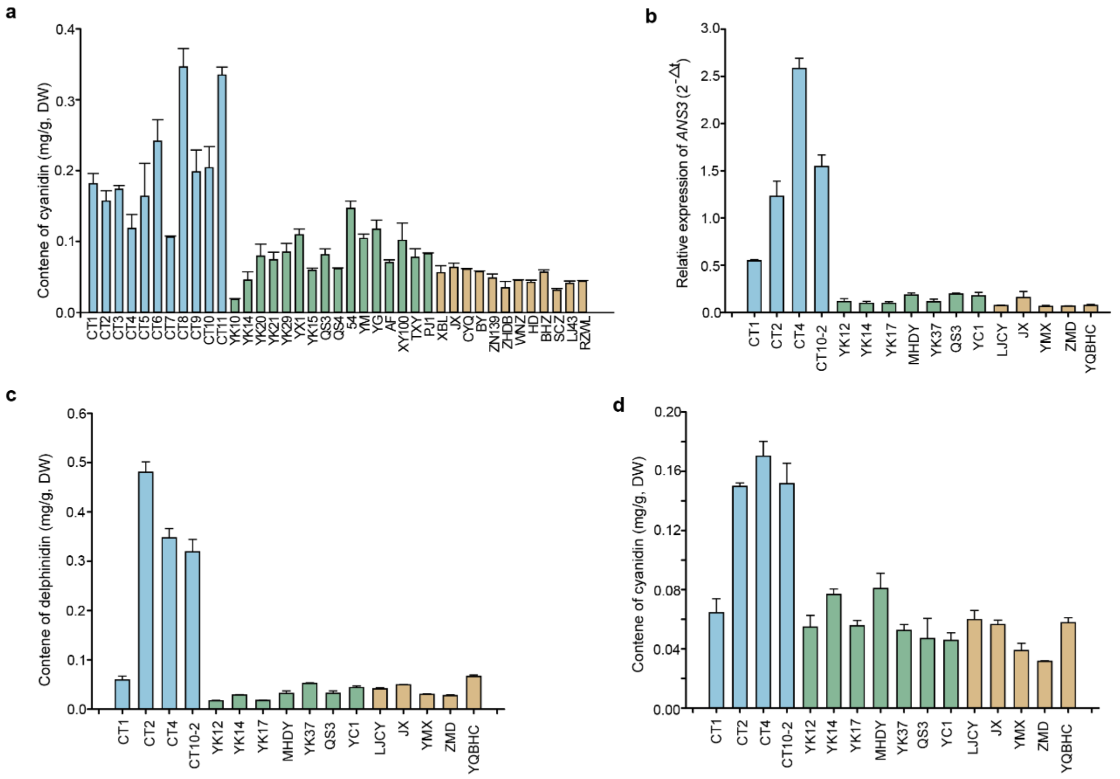


**Supplementary Fig. 7. Expression levels of ANS3 gene and anthocyanin content in different tea cultivars.** a) Cyanidin content in 39 tea plant accessions with three biological replicates per accession. Blue represents CT, green represents CSA, and yellow represents CSS. All 39 accessions were grown under the same environmental conditions and collected from Menghai, Yunnan. b) qRT-PCR analysis of *ANS3* gene expression in 19 tea plant accessions with three biological replicates per accession. All 19 accessions were grown under the same environmental conditions and collected from Pu'er, Yunnan. c,d) Content of delphinidin and cyanidin in 19 tea cultivars shown in panel b, with three biological replicates per accession.

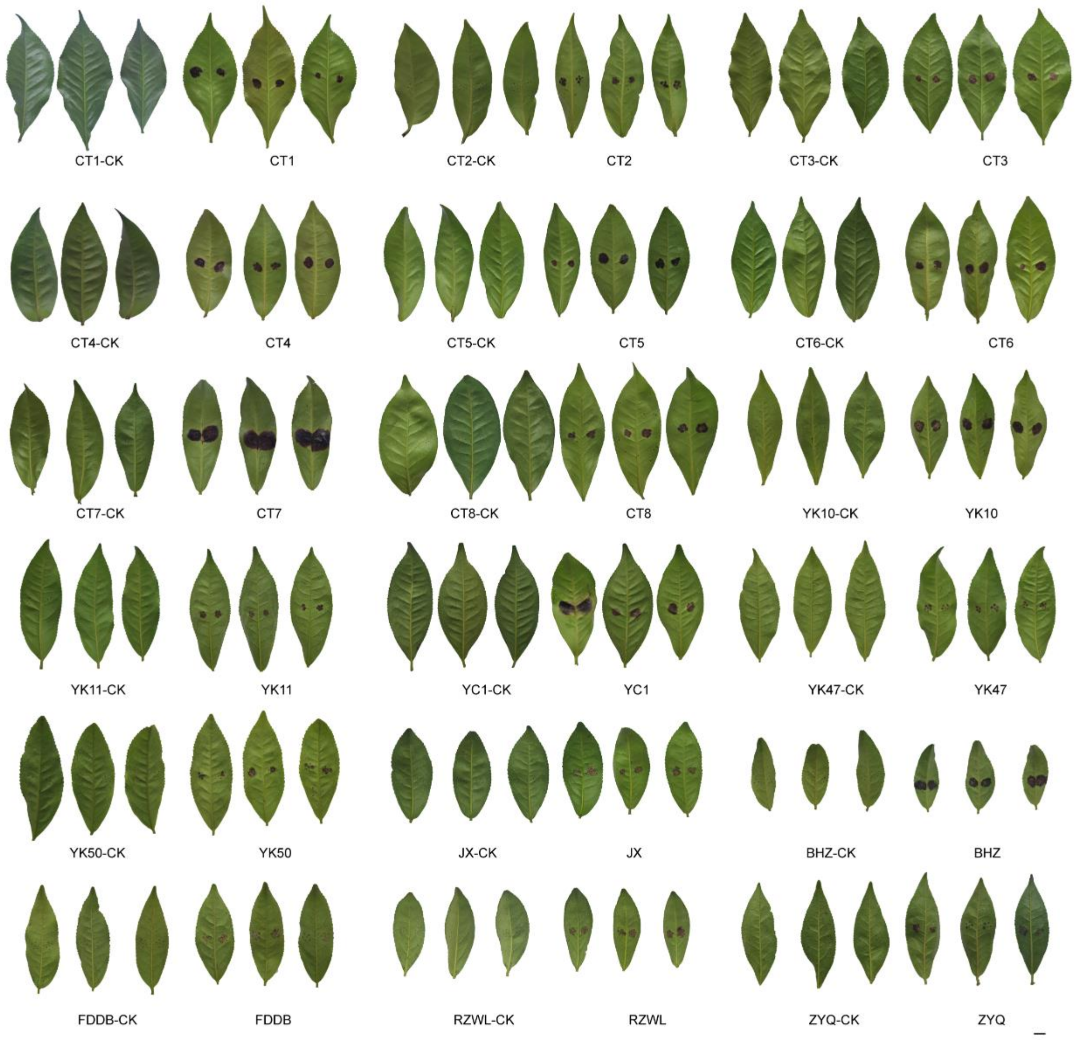


**Supplementary Fig. 8. Phenotypic characteristics of fungal infection in 18 tea plant accessions.** These accessions were grown under the same environmental conditions and collected from Menghai, Yunnan. Only 3 biological phenotypes were shown for each accession.

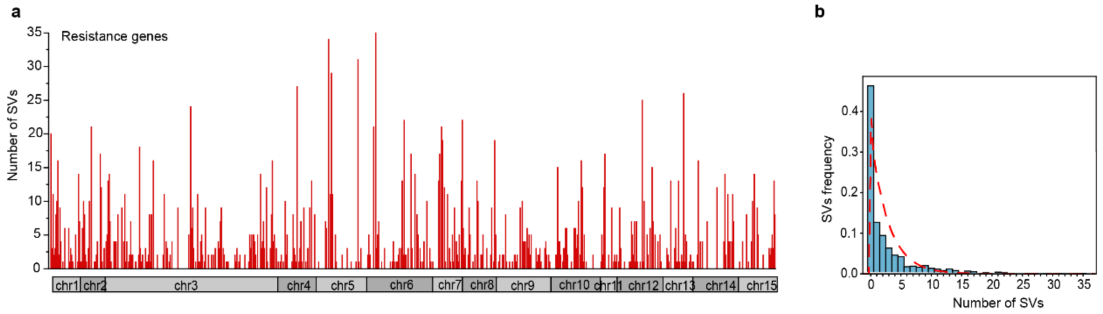


**Supplementary Fig. 9. Frequency of SVs occurring on R genes.** Genome-wide overlap between resistance genes and SVs. The red lines indicate the number of SVs detected for each gene. b) Shows the frequency of each SV.

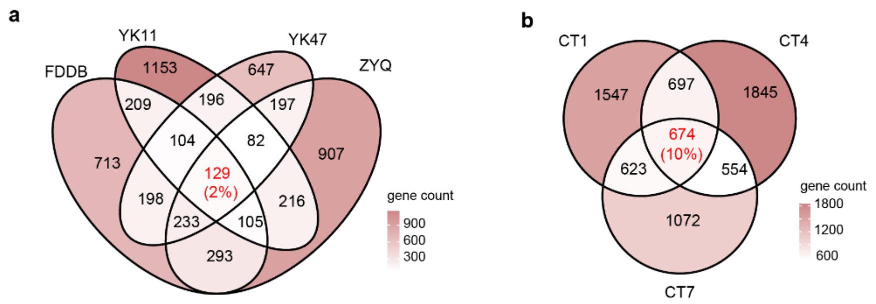


**Supplementary Fig. 10. Differentially expressed genes after *Colletotrichum. gloeosporioides* infection.** a) Differentially expressed genes in 4 cultivated accessions when comparing infected versus control. b) Differentially expressed genes in 3 CT accessions when comparing infected versus control.
